# A pan-generic marker panel for apples to enable genetic research and breeding across *Malus* species

**DOI:** 10.1101/2025.06.05.657940

**Authors:** Aafreen Sakina, Richard Tegtmeier, Hana Feulner, David Hickok, Anze Švara, Patrick Cho, Matthew Clark, Andrzej Kilian, Jeffrey Glaubitz, Qi Sun, Awais Khan

## Abstract

Wild *Malus* species harbor untapped genetic diversity to advance apple breeding, particularly for disease resistance and stress tolerance. However, existing marker panels, developed mainly using *Malus domestica* accessions, introduce ascertainment bias and limit detecting rare variants in wild species. We developed and validated a medium-density and cost-effective pan-generic 3K apple DArTag panel optimized to capture genome-wide variation across the *Malus* genus. The panel was constructed using conserved, syntenic, and collinear genomic blocks identified within the core genome of 13 *Malus* accessions for cross-species transferability. The panel was validated across three bi-parental mapping populations totaling 593 progeny. Across these populations, 2,461–3,234 SNP markers were polymorphic and 1,482–2,620 were informative. Each population contained over 900 multiallelic micro-haplotype loci, with several hundred loci exhibiting three or four distinct haplotypes. Markers were uniformly distributed across all 17 chromosomes, each containing between 60- 230 informative SNPs. The panel was further evaluated on 174 diverse germplasm accessions from 20 *Malus* species. It exhibited strong cross-species transferability, exceptionally low rates of missing data (<0.5%), and clear genetic differentiation between wild and domesticated accessions. Genome-wide association studies (GWAS) identified a major locus on chromosome 4 significantly linked to fruit length, weight, and width in addition to trait-specific associations on chromosomes 1, 6, 9, and 11. The cost-effectiveness of genotyping per sample (<$15), combined with these results, underscore the panel’s broad utility for quantitative trait locus (QTL) mapping in bi- parental and diverse populations, marker-assisted selection in the breeding programs, and genetic diversity analysis across the *Malus* genus.

## Introduction

Apple (*Malus domestica*) is one of the most important fruit crops grown in temperate regions worldwide (Duan et al. 2017). Apple breeding programs can greatly benefit from leveraging the extensive natural genetic diversity found in *Malus spp.* (Khan et al. 2021; Cornille et al. 2012). The wide genetic diversity found across *Malus spp*. is largely attributed to their predominantly outcrossing nature and the relatively weak domestication bottleneck (Dan et al. 2015). By broadening the genetic base beyond the limited number of elite cultivars currently in use, breeding programs can harness this diversity to enhance apple breeding (Migicovsky et al. 2021). Given the high heterozygosity, gametophytic self-incompatibility, prolonged juvenile phase, and linkage drag of unfavorable traits, a genomics-informed breeding approach can greatly improve apple breeding (Sabety et al. 2024).

Molecular markers are indispensable for genomics-assisted breeding and have been employed to develop genetic maps and advance the study of apple genetics (Longi et al. 2013). Previously, conventional molecular markers like Restriction Fragment Length Polymorphisms (RFLPs), Amplified Fragment Length Polymorphisms (AFLPs), Sequence Characterized Amplified Regions (SCARs), Simple Sequence Repeats (SSRs) and Diversity Arrays Technology (DArT) have been used to develop several saturated genetic maps (Hemmat et al. 1994; Conner et al. 1997; Maliepaard, 1998; Liebhard 2003; Fernández-fernández et al. 2008; Schouten et al. 2012). These maps have facilitated the genomic localization of economically important traits through many studies focused on novel QTL identification and marker-assisted selection (MAS) (Maric et al. 2010). However, these marker platforms are less suited for the fine dissection of functional genetic variations mainly due to their low density across the genome (Zhang et al. 2012).

Recent advances in genomic technologies and the availability of high-quality reference genomes in apples have revolutionized the study of genetic variations in apples (Chu et al. 2025, Khan et al. 2022; Sun et al. 2020; Švara et al. 2024a; Sabety et al. 2024). The development of SNP arrays such as Infinium® IRSC 8K and 20K apple SNP arrays, Expressed sequence tags (EST) based apple SNP genotyping platform, 50K SNP array and Affymetrix Axiom® Apple 480K SNP array has enabled the simultaneous genotyping of thousands of SNPs across the apple genome, and they have been widely used in the development of dense SNP-based linkage maps (Antanaviciute et al. 2012; Clark et al. 2014; Rymenants et al. 2020; Khan et al. 2012), QTL mapping (Falginella et al. 2015), pedigree-based analysis, association mapping (Kumar et al. 2013), and genomic selection (Kumar et al. 2012). However, the development of these arrays primarily relied on discovery panels consisting mainly of *M. domestica* scion cultivars (Rasheed et al, 2017). For example, the 8K SNP array was developed using 27 cultivars re-sequenced at low coverage, the 20K array was based on 14 major founders of European breeding programs, and the 480K array utilized data from 67 re-sequenced accessions (Chagne et al. 2012; Bianco et al. 2014; Bianco et al. 2016). This focus on domesticated apples could introduce ascertainment bias by failing to capture the rare and unique genetic variants present in wild *Malus* species (Peace et al. 2019). The main drawbacks of SNP arrays include a bias toward variants present in the accessions of the discovery panel used for the development of the array and under-representation of alleles with extreme allele frequencies (Giebel et al. 2021). Subsequently, the use of the Infinium 20K SNP array for genotyping 178

*Malus sylvestris* accessions revealed that 20% of the markers exhibited more than 10% missing values, 30% had a minor allele frequency (MAF) below 5%, and 6% were entirely monomorphic (Buiteveld et al. 2021). Wild apple species, which are reservoirs of novel alleles for disease resistance, abiotic stress tolerance, fruit quality, and rootstock traits, may harbor important genetic diversity that is not represented in domesticated varieties (Peace and Norelli 2009; Fazio et al. 2009, 2014; Duan et al. 2017; Norelli et al. 2017). Consequently, the SNP arrays may miss valuable genetic variation present in wild species, which can hinder efforts to breed apple cultivars with better traits. This underscores the importance of incorporating diverse reference panels including both domesticated and wild accessions to enable comprehensive and accurate genetic analyses (Yang et al., 2016).

The availability of high-quality draft genomes of domesticated apple and its wild relatives has provided a reference framework for developing novel genomic resources (<u>Malus all species | GDR</u> (rosaceae.org); Chu et al. 2025; Sabety et al. 2024; Švara et al. 2024a; Sun et al. 2020;). Advancements in next-generation sequencing (NGS) technologies offer the development of low-cost, high-throughput, and mid- density genetic marker screening platforms based on targeted genotyping-by- sequencing (GBS) (Endelman et al. 2024; Sabety et al. 2024). One such example of a targeted GBS marker platform is “DArTag” marker assays offered by Diversity Arrays Technology Ltd. (DArT; Canberra, Australia) (Hardigan et al. 2023; Endelman et al. 2024). DArTag marker panels utilize molecular inversion probe (MIP) genotyping to efficiently analyze large-scale single nucleotide polymorphisms (SNPs) and small indels. The custom-designed molecular probes amplify target regions containing SNPs, which are then sequenced using NGS. The sequenced amplicons are processed using DArT’s proprietary pipeline, enabling precise and comprehensive identification of SNPs (Sansaloni et al. 2011; Hardigan et al. 2023). Several DArTag panels have been developed for different crops such, as 2.5K and 4K DArTag panels for potato (*Solanum tuberosum L.*) (Endelman et al. 2024), 3K and 5K DArTag panels for cultivated strawberry (*F. × ananassa*) (Hardigan et al. 2023), 3K marker panel in cultivated blueberry (*Vaccinium spp*.), 3K marker panels in alfalfa (Zhao et al. 2023), 3K DArTag v.1 panel in cranberry (*Vaccinium macrocarpon*), 3K DArTag panel v.1 in cucumber (*Cucumis sativus)*, 3K DArTag panel v.2 for lettuce (*Latuca sativa*), DArTag panel v.1 in pecan (*Carya illinoinensis*), 3K DArTag panel v.1 for *Ipomoea batata* (https://breedinginsight.org/breeding-solutions/open-source-dartag-marker-panels). Twelve DArTag panels are also publicly accessible via CGIAR’s EiB platform (https://excellenceinbreeding.org/toolbox/services/mid-density-genotyping-service) to support genomic selection in breeding programs of CGIAR and its national partners.

Here, we report the development of a cost-effective 3K DArTag marker panel for *Malus*, designed using a diverse reference panel comprising both domesticated cultivars and wild accessions. The panel was validated in multiple bi-parental mapping populations and a *Malus* diversity panel. This high-throughput, transferable marker system enables robust QTL mapping, GWAS, linkage map construction, and population genomics analyses across diverse genetic backgrounds.

## Material and Methods

### 1. Development of DArTag marker panel

#### 1.1 Data preparation

Publicly available whole-genome assemblies of 13 *Malus* accessions, representing seven wild *Malus* species and five domesticated apples, were used for the development of the DArTag marker panel. Twelve assemblies were downloaded from the Genome Database for Rosaceae (GDR; https://www.rosaceae.org), while one, *M. orientalis* (Khan et al. unpublished), was obtained separately. The *M. domestica* assemblies included ‘Golden Delicious’ (GDDH13v1.1; Daccord et al. 2017), ‘Antonovka’ (Švara et al. 2023), ‘Gala’ (Sun et al. 2020), ‘Hanfu’ (Qin et al. 2023), and ‘Honeycrisp’ (Khan et al. 2022). The wild *Malus* species represented were *M. ioensis* (Švara et al. 2024a), *M. orientalis* (Khan et al. unpublished), *M. prunifolia* (Li et al. 2022), *M. sieversii* (Sun et al. 2020), *M. sylvestris* (Sun et al. 2020), *M. coronaria* (Švara et al. 2024a), *M. fusca* (Mansfeld et al. 2023), and *M. baccata* (Chen et al. 2019). These genomes were used for the core-genome identification and subsequent development of the DArTag marker panel.

#### 1.2 Syntenic core-genome construction

The assemblies were aligned to the reference genome GDDH13v1.1 (Daccord et al. 2017) using Minimap2 (version 2.17) (Li et al. 2018) with the -x asm10 alignment option to construct the syntenic core-genome. The parameter was employed to ensure a global alignment score greater than 400 and over 90% sequence identity with the reference genome for cross-species whole-genome alignment. The resulting BAM files were converted into BED format using a custom script to facilitate further analysis. Coverage depth per site was calculated using bedtools genomecov (Aaron et al. 2010), and coverage plots were generated by counting sites within sliding windows of 1 Mb. The data was curated and merged using Bedtools (Aaron et al. 2010) to refine the final output.

#### 1.3 Genus-wide variant calling

Illumina short-read sequences were obtained for 461 *Malus* accessions, as described in Liao et al. (2021). Sequences were downloaded from the Genome Sequence Archive (GSA) database (Genome Sequence Archive - CNCB-NGDC) under BioProject PRJCA004568, using the “fastq-dump” command from the SRA-Toolkit software (version 3.1.1). The sequence reads were aligned to the GDDH13 genome (Daccord et al. 2017) using BWA-MEM (v7.17) (Li and Durbin, 2009) with default parameters. The resulting alignments were then converted to BAM format using SAMtools (v1.16.1) (Li et al. 2009). Variants were called using the BCFtools package (v1.16) (Li et al. 2011), which produced SNP predictions from the GDDH13 alignments for all 461 *Malus* accessions. SNPs within the core genomic regions were filtered from the whole-genome sequencing data using the BCFtools package (Li et al. 2011). The SNPs were filtered into two categories: (1) high variance between wild and domestic apples, which are informative for hybrid populations; and (2) low F_st_ and high MAF between wild and domestic apples, which are informative for general apple populations. BCFtools (Li et al. 2011) was used to filter WGS high F_st_ SNPs by applying a series of criteria; biallelic variants were retained using the -m2 -M2 option; variants with more than 5% missing data were excluded across all samples using F_MISSING > 0.05. The data was filtered to ensure that no more than one individual in the wild population had missing data at a given site. Finally, the F_st_ values between wild and domesticated populations were calculated, and SNPs with a minimum F_st_ of 0.8% were retained to identify regions with significant population differentiation (min Q30; min MAF 1%; bi-allelic; maxMiss 5%; maxMiss wild 1; max F_st_WildDom0.8). To identify WGS lowF_st_ high MAF SNPs filter criteria were set to min Q30; min MAF 40 %; bi allelic; maxMiss 5%; maxMiss wild 1; max F_st_WildDom0.1. In addition, SNPs from the previously known 8K and 50K apple SNP arrays were filtered using criteria of a minimum quality score of Q30 and a MAF of at least 1% to ensure representation of variable SNPs within the core genome from these arrays. 128 QTL-linked SNPs from IRSCOA v1.0 apple SNP array were marked as “required” (Chagne et al. 2019). A total of 18,880 markers were submitted to DArT for QC, and 104 QTL-linked SNPs from IRSCOA v1.0 apple SNP array linked to known QTLs, which passed the design phase, were marked as “required” (Supplementary Table 1) (Chagne et al. 2019). The DArT algorithm selected SNPs in 54 base regions with high variant density for amplicon inserts and minimal flanking variant density for PCR primers. 9,134 markers (65.6%) passed DArT silico test for designing suitable flanking primers. Final marker selection was done among the “pass” SNPs for even distribution across genome-space and gene density. Ultimately, 3100 "genomic blocks" were selected for the final DArTag design. These regions are referred to as DArTag markers or amplicons, and each represents a specific genomic block targeted for amplification. For each of these 3,100 markers, a pair of primers (oligos) was designed to amplify a short DNA segment. Within each amplified region, multiple SNPs and micro-haplotypes are present. These SNPs and micro-haplotypes are the actual genetic variants used for genotyping, while the DArTag marker refers to the entire amplified region that contains them. This distinction is important throughout the manuscript; DArTag markers refer to the amplified genomic segments (not individual SNPs), each of which can harbor several informative variants used for downstream analyses.

### 2. Validation of DArTag marker panel

#### 2.1 Plant material

The marker panel was tested on three bi-parental mapping populations: Honeycrisp × Gala (n = 235), Gala × Demir (n = 179), and Gala × Chisel Jersey (n = 186), as well as on a diversity panel comprising 174 accessions representing a broad range of *Malus* species. These included *Malus angustifolia* (n= 10), *M. arnoldiana* (n =2), *M. baccata* (n=33), *M. coronaria* (n=5), *M. domestica* (n=5), *M.* ‘Evereste’ (n=1), *M. florentina* (n=1), *M. floribunda* (n=9), *M. hupehensis* (n=5), *M. hybrid* (n=2), *M. ioensis* (n=7), *M. micromalus* (n=12), *M. orientalis* (n=10), *M. fusca* (n=23), *M. prunifolia* (n=18), *M.* × *robusta* (n=10), *M. sargentii* (n=11), *M. sieversii* (n=2), *M. sylvestris* (n=5), and *M. zumi* (n=3). Leaf tissue for DNA extraction was collected from actively growing shoots, placed in 96-well plates, and shipped to Diversity Arrays Technology Pty Ltd (DArT, Canberra, Australia), where high-quality genomic DNA was isolated following their standard protocol (Jaccoud et al. 2001).

#### 2.2 Genotyping and allele calling pipeline

The Apple 3K DArTag marker panel is designed to call SNP variants from 54 bp amplicons, each strategically developed to include two or more SNPs within the target sequence. This design enables the identification of both individual SNPs and micro- haplotypes, which are contiguous sequence blocks within a single read containing multiple polymorphisms, including SNPs and small indels. The proprietary DArT genotyping platform provided a single SNP call per locus across the 3K marker panel, despite multiple SNPs within some amplicons (referred to as DArT markers onwards). To capture other variants and micro-haplotypes, raw amplicon reads were obtained from DArT and analyzed using a robust pipeline optimized for identifying multiple SNPs and micro-haplotypes within these amplicons (https://bitbucket.org/cornell_bioinformatics/amplicon/src/master/). The repository was downloaded from GIT (git clone https://bitbucket.org/cornell_bioinformatics/amplicon.git) (Zou et al. 2020). The pipeline de-multiplexes amplicons based on PCR primer sequences and subsequently calls novel alleles from FASTQ files by splitting reads using primer information. It identifies haplotypes across the population and optionally performs PCR error correction. Haplotype variants were obtained for each marker, and the results are compiled into a hap_genotype file, which is essentially a matrix with haplotype, genotype, and read count per allele information, and a haplotype allele fasta file which has haplotype sequence information for each allele. The haplotype file is converted to a haplotype VCF file using to_lep_map.pl script (https://bitbucket.org/cornell_bioinformatics/amplicon/src/master/), converts the haplotype genotype data into a haplotype VCF format. This script records four haplotype alleles per marker in each bi-parental mapping population as the pseudo standard nucleotide codes (A, C, G, T). Multiple variants within each amplicon were called by aligning the haplotype sequences generated by hap_genotype and the HaplotypeAllele.fasta file to the indexed reference genome GDDH13 (Daccord et al. 2017). The reference genome was indexed using bwa index. The resulting SNP VCF files were filtered using bcftools to retain sites with missing data < 30% and MAF > 0.01. The SNPs and haplotypes identified are referred to as “new pipeline SNP and micro-haplotype markers” in this paper.

#### 2.3 Bi-parental mapping population

The marker panel was tested with three bi-parental mapping populations: Honeycrisp × Gala (n = 235), Gala × Demir (n = 179), and Gala × Chisel Jersey (n = 186). Principal component analysis (PCA) was performed to validate the robustness of the panel in assessing genetic variation among parental replicates and F_1_ individuals. PCA was performed separately using SNP and micro-haplotype data from three different bi-parental mapping populations. The VCF file was imported into R version 4.3.2 using the vcfR package (Knaus and Grunwald, 2017) and converted into a genlight object with the vcfR2genlight function (Knaus and Grunwald, 2017). PCA was conducted on the genlight object using the glPca function from the adegenet package, retaining the first 10 principal components (PCs) (Jombart, 2008). The PCA results were visualized using ggplot2 (Wickham, 2016), with parents and F_1_s colored distinctly.

#### 2.4 SNP filtering and informative marker selection

SNP VCF files were first filtered to retain only those variants with consistent genotypes across biological replicates of both parents. Specifically, markers were retained if each parent exhibited identical genotype calls across all replicates and had no missing genotype values. Among the consistent markers, monomorphic loci were identified as those where both parents shared the same homozygous genotype and subsequently removed. The remaining polymorphic markers were then screened for informativeness. Markers were retained if they met one of the following conditions: 1. One parent was homozygous, and the other was heterozygous, 2. both parents were heterozygous. The resulting informative marker dataset was used to plot the chromosomal distribution of informative markers across the 17 chromosomes. Marker counts per chromosome were calculated and visualized using a bar plot to evaluate genome-wide coverage of polymorphic markers.

#### 2.5 Diversity population

The marker panel was evaluated on a diverse *Malus* population comprising 174 unique genotypes and 247 total samples, including technical replicates to ensure genotyping consistency. This population represented a broad spectrum of *Malus* diversity, encompassing 20 species with varying sample sizes (Supplementary Table 2).

#### 2.6 Genetic properties of populations

##### 2.6.1 Principal component analysis

PCA was conducted using a total of 15,885 SNP markers (from the new pipeline), which were previously filtered to remove insertions and deletions (INDELs) using TASSEL v5 (Bradbury et al. 2007). The resulting VCF file was imported into R (version 4.3.2) using the vcfR package (Knaus and Grunwald, 2017). SNP data were read using the read.vcfR function and subsequently converted into a genlight object *via* the vcfR2genlight function from the adegenet package (Jombart 2008) for multivariate analysis. PCA was performed using the glPca function (Jombart 2008), retaining the first two PCs for downstream visualization and interpretation. The PCA results were visualized using the ggplot2 package (Wickham, 2016). Samples were color-coded by species, with distinct species-specific palettes generated using the randomcoloR packages (Ammar 2006).

##### 2.6.2 Phylogenetic analysis

To investigate genetic relationships among the *Malus* accessions, a phylogenetic tree was constructed using SNP data (from the new pipeline) in R (version 4.3.2). 16,002 SNPs were used, and missing data were handled by excluding loci with over 50% missing values. The cleaned matrix was converted into a numeric matrix and subsequently into a genlight object using the dartR package (Gruber et al. 2018), with ploidy set to 2. Monomorphic and all-missing loci were filtered out using gl.filter.monomorphs and gl.filter.allna. A neighbor-joining (NJ) tree was computed based on bitwise genetic distances using the bitwise.dist function from poppr (Kamvar et al. 2015), and the NJ algorithm implemented in the ape package (Paradis et al. 2004). Bootstrap support values were estimated using 1,000 replicates with a custom bootstrap function that resamples SNP columns and reconstructs NJ trees. Node support values were assigned using the boot.phylo function. The final NJ tree with bootstrap support was visualized in R and saved in Newick format using write.tree from the ape package (Paradis et al. 2004). The Newick file was then imported into the Interactive Tree of Life (iTOL) platform (https://itol.embl.de/) for enhanced visualization, including taxon-specific color coding and annotations (Letunic and Bork, 2021).

##### 2.6.3 Population structure analysis

The population genetic structure of *Malus* accessions was assessed using the model- based Bayesian clustering algorithm implemented in STRUCTURE v2.3.4 (Pritchard et al. 2000). This method uses a Markov Chain Monte Carlo (MCMC) approach to estimate the proportion of each individual’s genome derived from *K* ancestral populations, aiming to minimize deviations from Hardy–Weinberg equilibrium and linkage disequilibrium within inferred clusters. Analyses were conducted using the admixture model with correlated allele frequencies. Five independent replicates were run for each value of *K* (ranging from 2 to 12), each with a burn-in period of 10,000 iterations followed by 50,000 MCMC iterations to ensure convergence and model stability. The results were processed using CLUMPP v1.1.2 to align cluster assignments across replicate runs and identify consistent clustering patterns (Jakobsson and Rosenberg 2007). The aligned outputs were also visualized using DISTRUCT v1.1 to display individual admixture proportions (Rosenberg, 2004). Final plots and summary statistics were generated using the Pophelper R package (Francis, 2017), facilitating visualization and interpretation of population structure across varying *K* values (Francis et al. 2017).

#### 2.7 LD decay

Linkage disequilibrium (LD) decay across the genome was analyzed using PLINK v1.9 (Chang et al. 2015). SNPs were filtered to retain variants with a MAF ≥ 0.01, a missing genotype rate per SNP ≤ 10%, and a missing genotype rate per individual ≤ 50%. LD decay was computed using pairwise squared correlation coefficients (r²) within a 50 kb sliding window across all chromosomes. The following PLINK options were used to define the LD window: --ld-window 99999: no limit on the number of variant pairs considered, --ld-window-kb 50: restricts comparisons to variant pairs within 50 kilobases, --ld-window-r2 0: includes all SNP pairs regardless of their r² value. The final dataset was then converted to PLINK binary format (.bed, .bim, .fam) using --make-bed for downstream analyses. The LD decay pattern was visualized in R (v4.3.2) using the ggplot2 package (Wickham, 2016). The absolute genomic distance between SNP pairs was computed, and LOESS smoothing was applied to estimate the decay of LD (r²) with increasing physical distance. The half-LD decay distance was calculated as the physical distance at which the fitted LOESS curve dropped to half the maximum observed r² value. Multiple visualizations were generated using raw and smoothed LD values to represent decay patterns at different resolutions (base pairs and kilobases).

#### 2.8 Marker distribution and coverage

The SNP marker distribution across chromosomes was evaluated using the .bim file generated by PLINK v1.9 (Chang et al. 2015). SNP positions were binned into 1 Mb windows, and the number of markers per bin was summarized using dplyr and tidyr (Wickham et al. 2023; Wickham et al. 2024). SNP coverage was displayed as point plots, showing exact marker locations and densities across chromosomes. A chromosome-wide summary of LD was performed by grouping pairwise SNP comparisons based on chromosome identity.

#### 2.9 Genome wide association study

Phenotypic data for fruit size traits were obtained from the publicly accessible USDA National Plant Germplasm System (NPGS) *via* the GRIN-Global database (https://npgsweb.ars-grin.gov/gringlobal/search?q=APPLE). Three quantitative fruit traits were analyzed; fruit length, fruit width, and fruit weight for 169 accessions. Measurements were recorded as part of standard germplasm evaluation protocols. Fruit length and *fruit width* were recorded as the average of 10 mature fruits collected from a vigorous tree, measured in millimeters (mm). Fruit weight was expressed as the mean weight (in grams) of 10 mature fruits at physiological maturity. The SNP VCF file was filtered to retain only SNPs by removing indels using bcftools (Li et al. 2011). GWAS was conducted for fruit quality traits using the General Linear Model (GLM) and the Mixed Linear Model (MLM), both accounting for population structure using the first five PCs and Q matrix. The MLM additionally accounted for kinship by incorporating (K), a kinship matrix, accounting for relatedness. Analysis was performed using the Genomic Association and Prediction Integrated Tool (GAPIT) package version 3.5.0 (Wang et al. 2021).

## Results

### 1. Development of DArTag marker panel

#### 1.1 Core genome construction

Thirteen *Malus* genomes representing phylogenetically and geographically distinct species were selected to construct the core genome. These assemblies collectively span the genetic diversity across the *Malus* genus and form the foundation for identifying conserved genomic regions in the core genome. While the ideal approach would involve designing markers exclusively from genomic regions shared across all 13 genomes, this was not feasible. In practice, only 2.1% of the reference genome, primarily encompassing 80.1% of annotated protein-coding gene regions, was conserved across all assemblies. A hierarchical strategy was employed to address potential gaps in marker coverage. Markers were first placed in regions present in ≥12 genomes, ensuring a high-confidence representation of conserved loci. Subsequently, remaining gaps were filled using regions found in ≥8 genomes, maximizing genomic coverage while maintaining cross-genome relevance. This approach resulted in marker coverage of approximately 35 Mb out of the 625 Mb reference genome. Among the selected genomes, Hanfu and Honeycrisp exhibited the highest genome coverage, each with 99.50%, followed closely by *M. sieversii* (99.30%), Gala (99.20%), and *M. sylvestris* (98.50%). Antonovka also showed high coverage at 98.40%. Wild North American species such as *M. fusca* (95.80%), *M. orientalis* (95.10%), and *M. coronaria* (94.00%) demonstrated moderately high coverage, while *M. ioensis* had a slightly lower value at 92.60%. *M. baccata* and *M. prunifolia* showed the lowest genome coverage among the assemblies, with 89.60% and 84.60%, respectively.

#### 1.2 Reference genome coverage across multiple assemblies

To evaluate the extent of conserved genomic regions across *Malus* assemblies, we calculated reference genome coverage based on presence across 12, 11, and 10 assemblies, respectively (Table 1). When considering regions conserved in ≥12 assemblies, only 2.1% of the 625 Mb reference genome was covered, corresponding to 13.2 Mb of sequence. These regions, however, captured 80.1% of annotated protein-coding gene space, indicating a strong representation of functionally important loci. Chromosome-level coverage ranged from 0.7% (Chr01) to 2.7% (Chr07 and Chr10), with the highest average marker density observed on Chr04 (377 bp) and Chr14 (397 bp) and no coverage on Chr00 (unplaced contig). Expanding the inclusion threshold to regions conserved in 11 assemblies resulted in 5.6-fold greater coverage, spanning 34.8 Mb or 5.6% of the reference genome. This set included 76.5% of protein-coding gene regions, suggesting a modest trade-off between broader genomic coverage and gene content. Per-chromosome coverage increased substantially from 2.9% (Chr01) to 6.8% (Chr15). Fragment spacing also increased slightly, averaging between 431 bp (Chr00) and 522 bp (Chr13). Further relaxing the threshold to include regions present in ≥10 assemblies yielded the most extensive coverage: 55.0 Mb, representing 8.8% of the reference genome. However, the proportion of gene space captured dropped to 69.1%, reflecting the inclusion of more variable, non-genic regions. Chromosome-level coverage reached up to 12.7% (Chr15), with average fragment sizes ranging from 534 bp (Chr01) to 628 bp (Chr06) (Figure 1).

**Figure 1.**
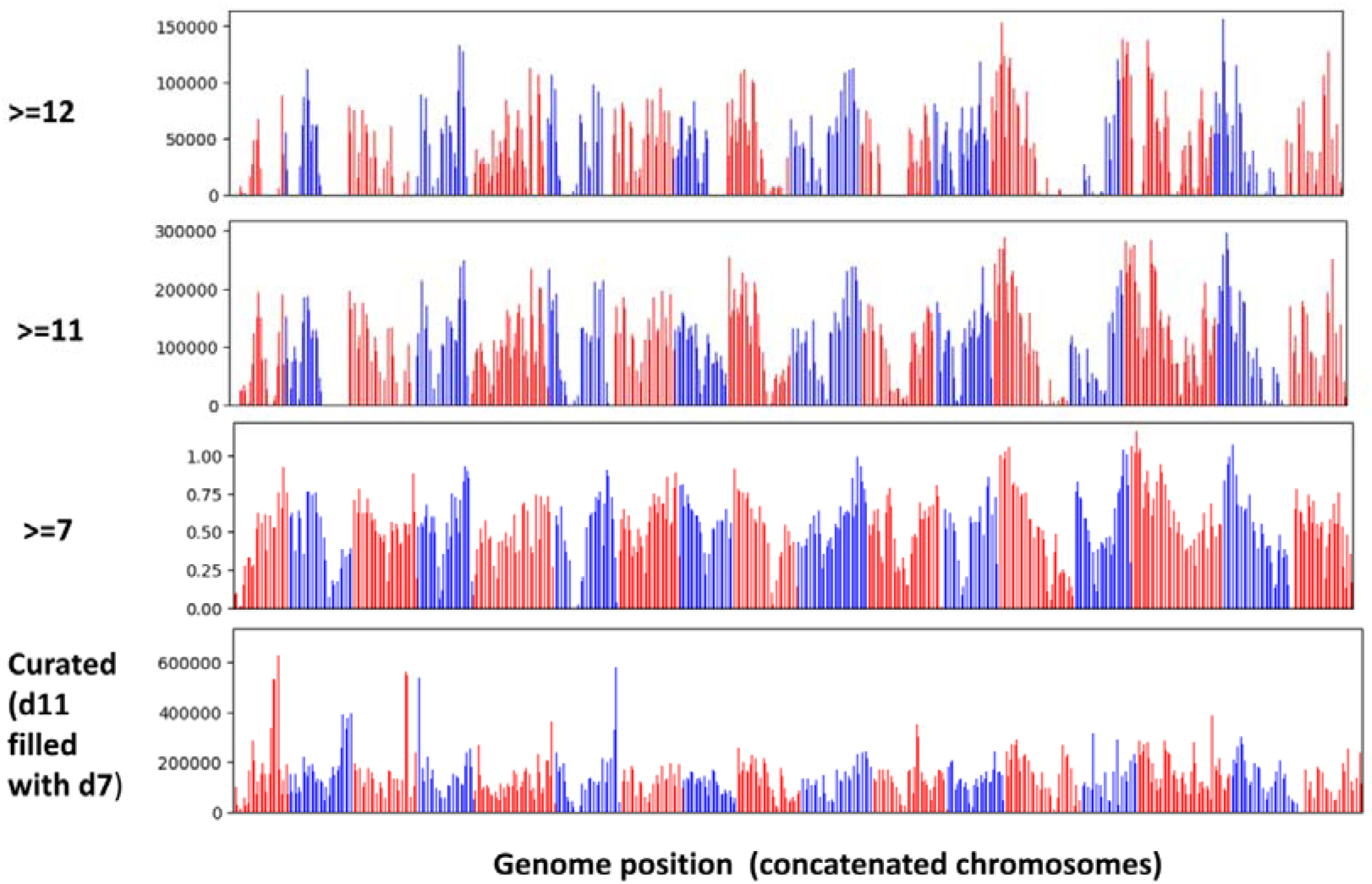
Genomic distribution of conserved regions across multiple *Malus* genome assemblies used to design DArTag marker panel. Bar plots show the genomic distribution of regions conserved across *Malus* assemblies using three thresholds: regions present in ≥12 assemblies, ≥11 assemblies, and ≥7 assemblies, respectively. The bottom panel represents a curated marker panel, where regions present in ≥11 assemblies were supplemented with regions from ≥7 assemblies to fill gaps. Red and blue bar denote alternating chromosomes for visual clarity.

**Table 1.**
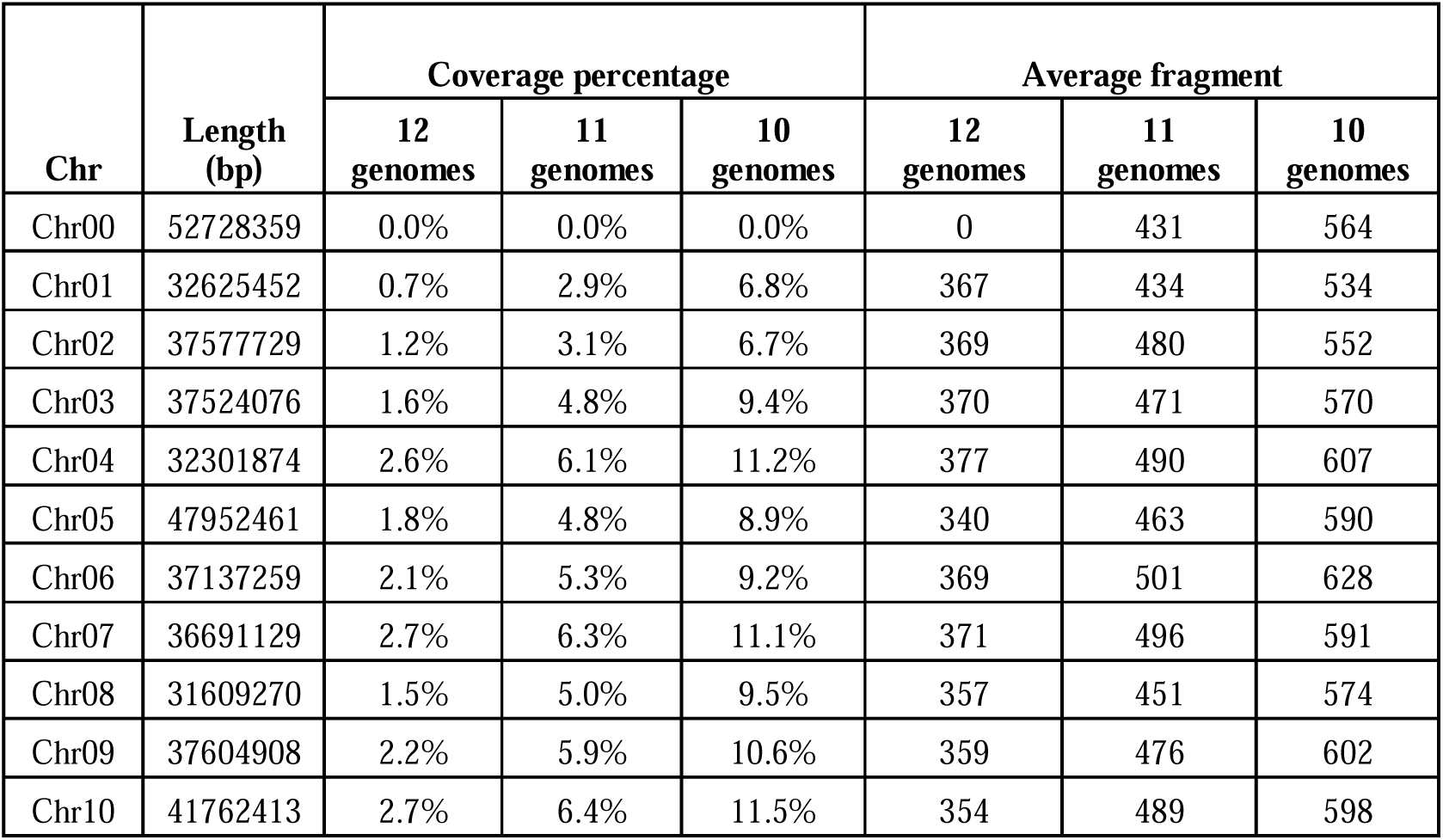
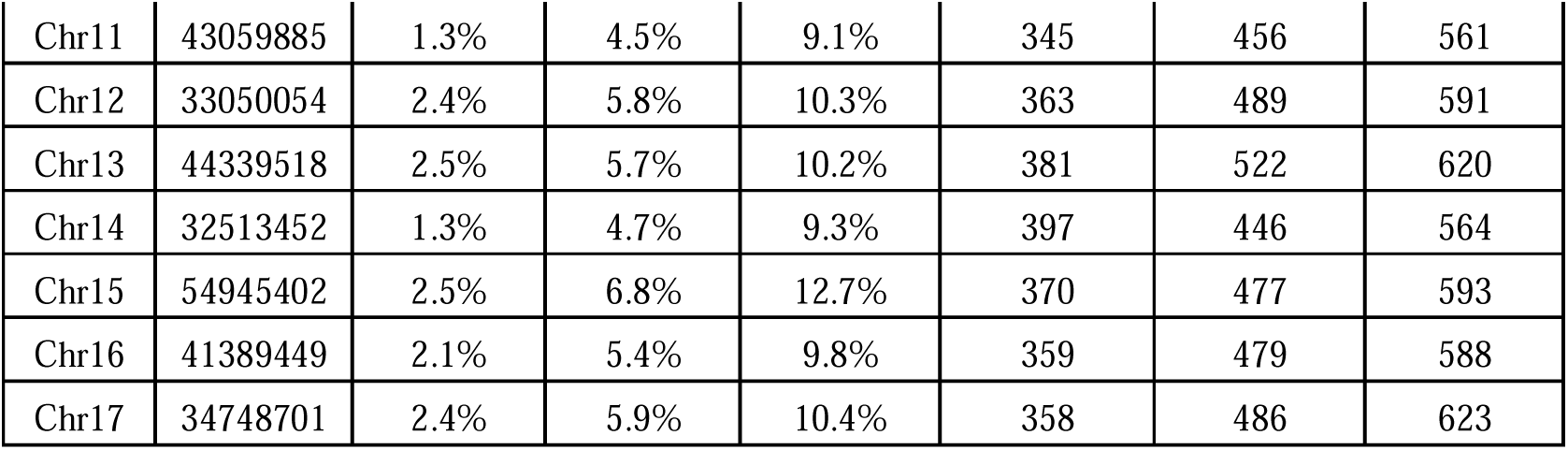
Chromosome-wise reference genome coverage by conserved regions across 12 *Malus* genome assemblies. This table summarizes the extent of conserved genomic regions mapped onto each chromosome of the reference genome. “Length” refers to the total physical length (in base pairs) of each chromosome. “Coverage percentage (12 genomes),” “Coverage percentage (11 genomes),” and “Coverage percentage (10 genomes)” indicate the proportion of each chromosome covered by regions conserved across all 12 genomes, at least 11 genomes, and at least 10 genomes, respectively. “Average fragment” values represent the average length (in base pairs) of continuous conserved regions detected at each coverage threshold.

#### 1.3 Marker filtering and DArTag panel design

The DArTag marker panel was developed through a two-phase marker selection process designed to optimize genomic coverage. In phase one, the DArT algorithm was employed to identify SNP-rich 54 bp regions suitable for amplicon design, while minimizing flanking sequence variation to ensure primer efficiency. A total of 13,919 candidate SNP loci were submitted for *in silico* evaluation. Additionally, 104 SNPs from the IRSCOA v1.0 apple SNP array, each linked to known QTLs, were prioritized for inclusion based on their association with key disease resistance and fruit quality traits (Chagné et al. 2019). These include major resistance loci for apple scab (*Rvi6*, *Rvi11*, *Rvi2*, *Rvi4*, *Rvi15/Vr2*, *Rvi3*, *Rvi12*), fire blight (*RLP1*, *MR5*, *FBE*), woolly apple aphid (*Ermis*), and powdery mildew (*Pl2*). SNPs related to fruit quality traits represented by associated genes include fructose content (*LG1Fru*), esters (*MdAAT1*), acidity (*Acidity-LG8*, *Ma1*), red skin color (*MYB10*), fruit firmness (*MdPG1*, *LG15-BB*), vitamin C (*GGP3*), ethylene (*MdACO1*, *MdACS1*), bitter pit (*LAR/Ma*, *Bp13*), polyphenols and bitter pit (*LAR1*), and additional traits such as type 2 red flesh (*MYB110*) and maturity time (*Maturity-LG3*) were also included (Chagné et al. 2019). Of all submitted markers, 9,134 (65.6%) passed the *in silico* design filters. In phase two, marker selection was refined to ensure even representation across the genome based on gene density. From the passing markers, 3,100 genomic blocks, each representing approximately equal numbers of genes, were selected to ensure balanced coverage. All 104 QTL-linked SNPs were retained in the final panel.

Table 2 summarizes the filtering statistics from different SNP sources used in panel development. Of particular interest were the 13,073 high F_st_ SNPs and 3,105 low F_st_, high MAF SNPs, which had sufficient flanking SNP support (≥3 and ≥2 flanking SNPs, respectively) and passed conserved primer design checks. These sets are visualized in Figure 2, displaying their genomic density across the *Malus* reference genome. The total number of SNPs in the final design is described in Table 3.

**Figure 2.**
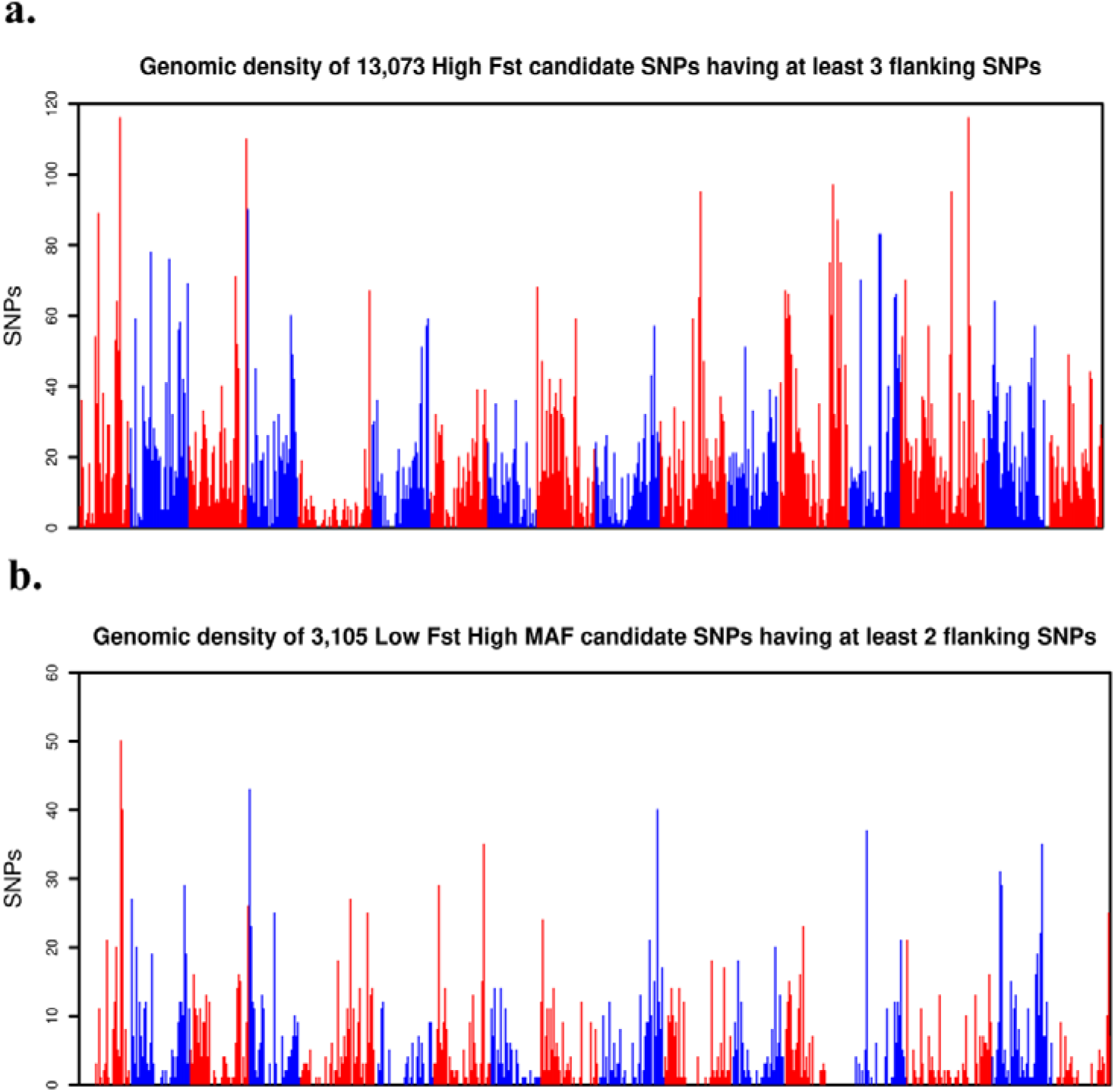
Genomic distribution of candidate SNPs across the apple genome a) Genomic density of 13,073 high- F_st_ candidate SNPs, each flanked by at least three neighboring SNPs, indicating genomic regions with strong population differentiation. b) Genomic density of 3,105 low- F_st_, high-MAF candidate SNPs, each supported by at least two flanking SNPs, selected for their broad allele representation across diverse apple accessions. The x-axis represents genomic positions (in megabases, Mb), concatenated across all chromosomes of the apple reference genome. The y-axis shows the number of candidate SNPs within each genomic window, reflecting local marker density and distribution. Alternating colors along the x-axis indicate transitions between chromosomes, allowing chromosome boundaries to be visually distinguished across the concatenated genome.

**Table 2.** Summary of SNP marker selection and filtering across different sources for DArTag panel development during phase 1. The table outlines the number of SNPs at each stage of the filtering pipeline across five sources: whole-genome sequencing (WGS) high- F_st_ and low- F_st_, high-MAF candidates, the 9K and 50K apple arrays, and a curated set of 128 QTL-linked SNPs.

| SNP Source | WGS High $F_{st}$ | WGS Low $F_{st}$ HighMAF | 9K | 50K | 128 QTL-linked |
| --- | --- | --- | --- | --- | --- |
| Total original SNPs | 82,310,931 | 82,310,931 | 7,399 | 52,638 | 114 |
| Pass Filter | 60, 291 | 14,596 | 1,056 | 6,057 | 104 |
| minFlankSNPs | 3 | 2 | 2 | 2 | 0 |
| conservedPrimers | 13,073 | 3,105 | 361 | 2,238 | NA |
| ReadyForDArT | 13,073 | 3,105 | 361 | 2,238 | 104 |
| Total ready for DArT | 18,880 |  |  |  |  |

**Table 3.**
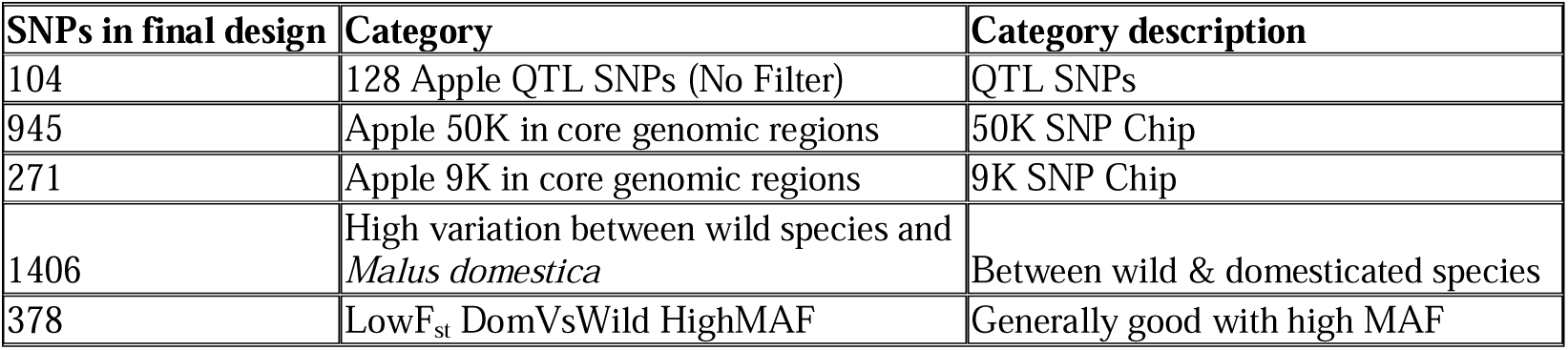
Classification of SNPs included in the final DArTag panel design. This table summarizes the composition of the final SNP set (n = 3,100), categorized by source and selection criteria. The panel integrates 104 QTL-linked SNPs without filtering, SNPs from legacy apple arrays (9K and 50K) located in core genomic regions, and SNPs chosen based on population differentiation and allele frequency metrics.

### 2. Validation of DArTag marker panel

#### 2.1 Bi-parental F_1_ mapping populations

The newly developed DArTag marker panel was validated on the Honeycrisp × Gala bi-parental population, consisting of 235 individuals (Figure 3). A total of 3,320 SNPs obtained from the new SNP pipeline were analyzed. Among these, 3,137 were polymorphic and 2,841 were found to be informative, while 183 markers showed genotype inconsistencies between parental replicates. The genomic distribution of SNPs revealed even coverage across the 17 apple chromosomes. Chromosome-wise marker distribution of informative markers ranged from 112 markers on Chr13 to 230 markers on Chr15, with the remaining chromosomes containing between 115 and 212 markers. Among 3,097 haplotype markers, 829 loci were identified as multiallelic. Specifically, 612 loci exhibited three haplotypes, and 217 loci displayed four distinct haplotypes, indicating a high level of allelic diversity. The DArTag markers included 3,099 SNPs, with 13 markers differing between replicates. Of these, 1,676 SNPs were polymorphic and 1,482 were informative.

**Figure 3.**
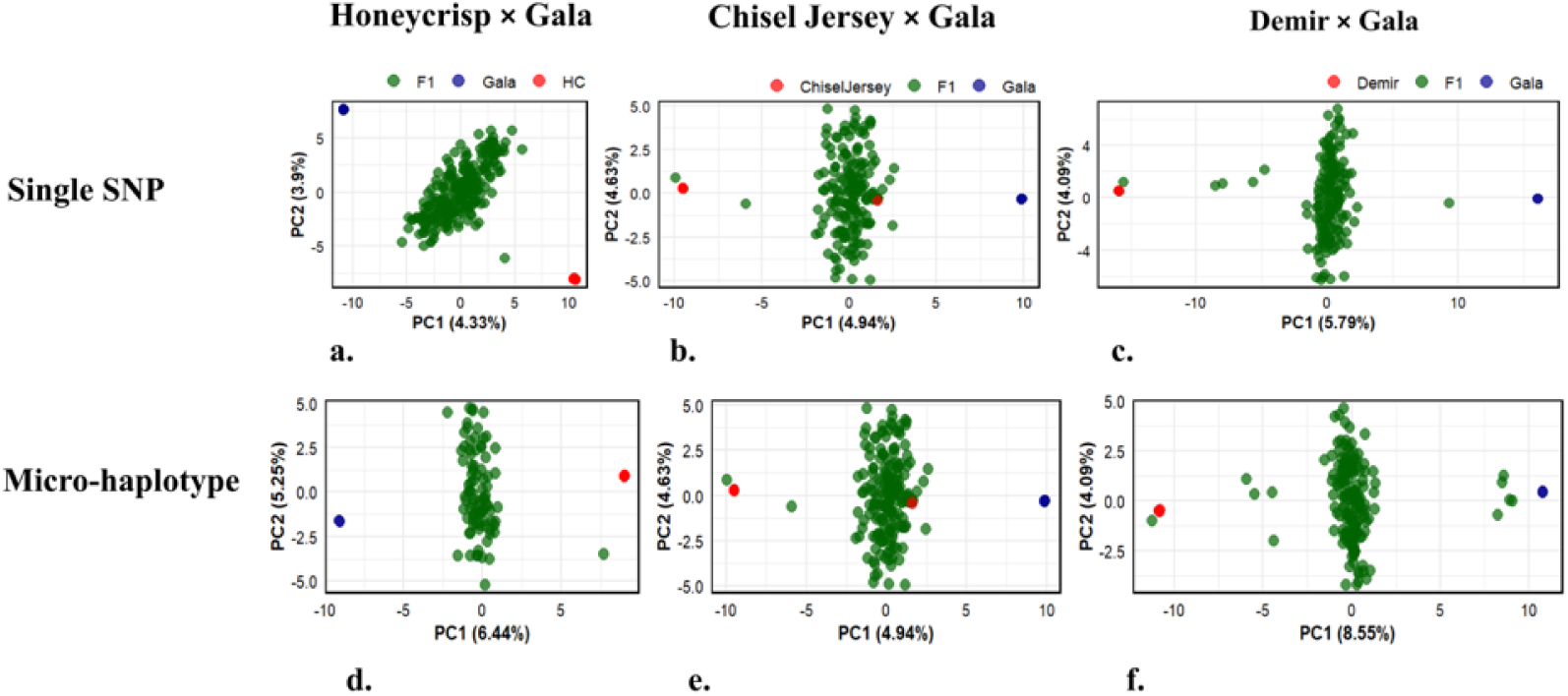
Principal component analysis of three bi-parental mapping apple populations based on new pipeline SNP (upper panel) and micro-haplotype markers (lower panel) (a and d) Honeycrisp × Gala population (n = 235), (b and e) Chisel Jersey × Gala population (n = 186), and (c and f) Demir × Gala population (n = 172). Each dot represents an individual. F_1_ progeny are shown in green, while parental genotypes are colored separately: red for Honeycrisp (a, d), Chisel Jersey (b, c), and Demir (e, f); and blue for Gala.

The Chisel Jersey × Gala bi-parental population (n = 186) was genotyped using the SNP panel comprising 3,097 markers (Figure 3). A total of 3,099 DArTag markers were initially analyzed across 186 progeny. Of these, 11 markers showed genotype discrepancies between technical parental replicates, and 1,824 markers were polymorphic, with 1,560 classified as informative. Using the refined set of 3,097 markers from the new pipeline, 171 were found to differ between parental replicates. The refined dataset yielded 2,461 polymorphic and 1,941 informative markers. A total of 1,513 informative SNPs from this population were successfully placed across the 17 apple chromosomes. Chromosome 7 contained the highest number of markers (132), while Chromosome 1 had the fewest (60). Marker counts for the remaining chromosomes ranged from 60 to 130. Among these, 904 loci were identified as multiallelic and subsequently excluded from PCA. Haplotype-based analysis revealed that, of the multiallelic loci, 632 loci exhibited three distinct micro-haplotypes, while 272 loci had four micro-haplotypes, indicating high allelic diversity in the panel. Notably, one parental sample (Chisel Jersey_002) appeared to cluster incorrectly with progeny, suggesting potential sample mislabeling or contamination.

The Demir × Gala bi-parental population was evaluated using both the pipeline developed by Diversity Arrays Technology (DArT) and new SNP pipelines (Figure 3). The newly developed SNP panel included 3,514 markers, of which seven loci exhibited more than two alleles and were excluded from further biallelic analysis. The population consisted of 172 individuals. Haplotype analysis using a set of 3,097 markers revealed 940 multiallelic loci. Of these, 680 loci exhibited three distinct haplotypes, and 260 loci exhibited four haplotypes, reflecting substantial haplotypic diversity in the panel. In the DArT panel pipeline, 3,099 SNP markers were used on a population of 185 individuals. Among these, 30 markers showed discrepancies between replicates, 1,726 were polymorphic, and 1,336 were informative. The updated pipeline showed significant improvements: among the 3,514 markers evaluated, only 111 exhibited genotype differences between technical replicates. Furthermore, 3,234 markers were polymorphic and 2,620 were informative. A total of 2,407 informative markers were successfully mapped across the 17 apple chromosomes. Chromosome 15 had the highest number of markers (214), while Chromosome 13 had the fewest (94). The remaining chromosomes had marker counts ranging from 102 to 173.

#### 2.2 Genetic properties of diversity panel

##### 2.2.1 Principal component analysis

PCA was performed on SNP genotype data from multiple *Malus* species to assess genetic relationships. The first two PCs, PC1 and PC2, explained 23.23% and 17.37% of the total genetic variance, respectively, and were used to visualize genetic clustering among accessions (Figure 4). Species-level grouping was supported by 95% confidence ellipses, which delineate the distribution of each species in PC space. North American species *M. angustifolia*, *M. ioensis*, and *M. coronaria* formed a tight cluster, distinct from *M. fusca*, indicating divergent genetic backgrounds. Interestingly, two accessions of *M. fusca* clustered apart from the main *M. fusca* group, while *M. florentina* was positioned closely to *M. fusca*, suggesting potential genetic relatedness. One accession of *M. ioensis* also clustered separately, highlighting intraspecific divergence. European and Central Asian wild species *M. sieversii*, *M. orientalis*, and *M. sylvestris* clustered near *M. domestica*, reflecting their known contributions to the domesticated apple gene pool. Several accessions of *M. robusta* and *M. sargentii* occupied intermediate positions or showed partial overlap with *M. baccata*, potentially indicating shared ancestry or historical admixture events. Overall, the clustering pattern underscores clear genetic differentiation between North American and Eurasian wild *Malus* species and supports evidence of divergence and gene flow among wild and cultivated lineages (Figure 4).

**Figure 4.**
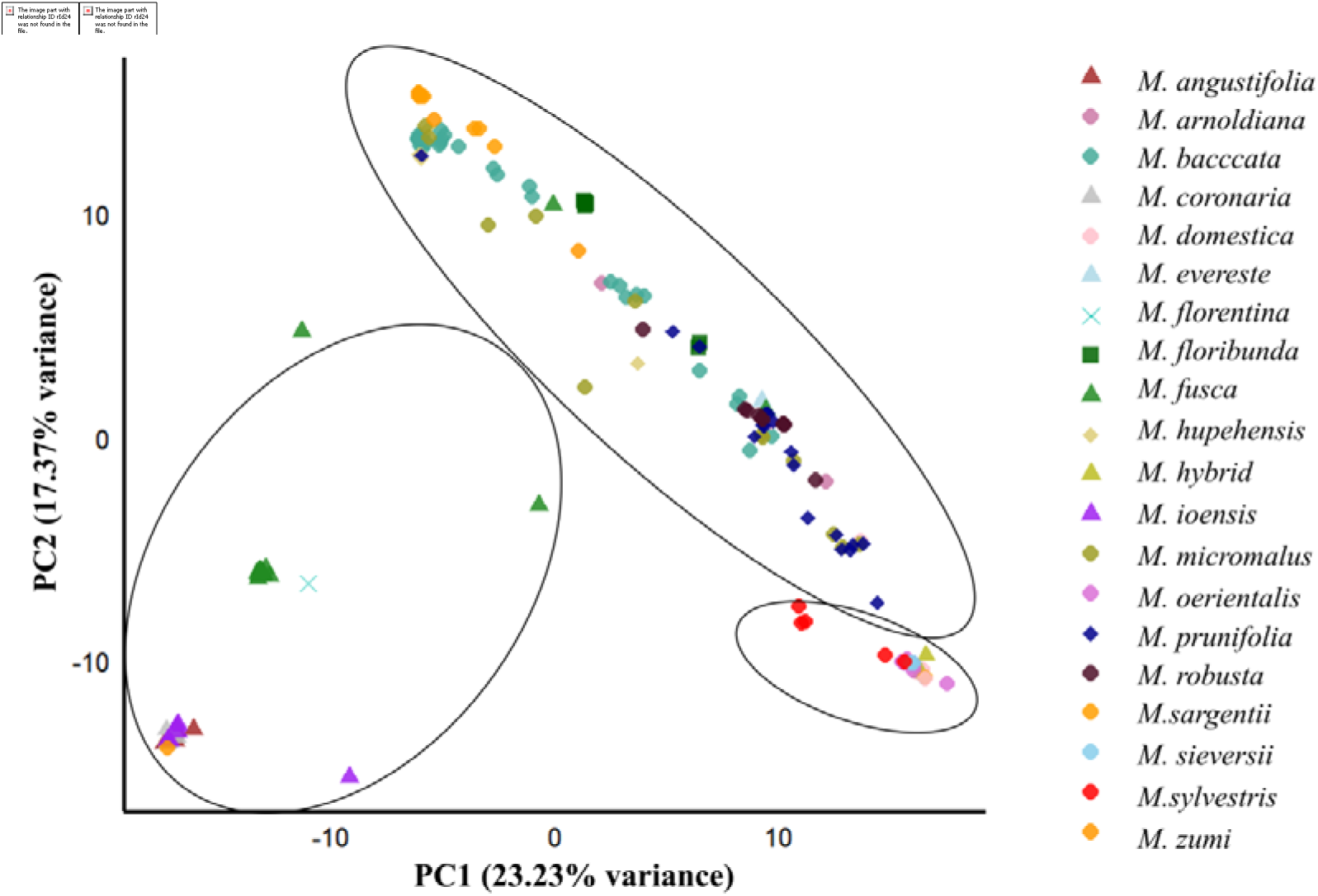
PCA of diverse *Malus* species based on genome-wide SNP data. Each point represents an individual accession, colored and shaped by species identity. The first two principal components (PC1 and PC2) explain 23.235 and 17.37% of the total genetic variance, respectively, and capture the major axes of genetic variation. Distinct clustering is observed for North American (*M. angustifolia*, *M. ioensis*, *M. coronaria, M. fusca*), Eurasian species (*M. sieversii*, *M. orientalis*, *M. sylvestris*) and East Asian *Malus* species. To improve visual distinction among taxa, selected species including *M. angustifolia*, *M. fusca*, *M. coronaria*, and *M. ioensis* were assigned unique point shapes (triangles), while remaining species were displayed using standard plotting symbols such as circles or squares. Solid outlines represent 95% confidence ellipses for each species, indicating the expected distribution of accessions around the species centroid in PC space.

##### 2.2.2 Phylogenetic analysis

A maximum likelihood phylogenetic tree was constructed to infer the evolutionary relationships among diverse *Malus* accessions (Figure 5). The tree revealed distinct clades corresponding to species-level groupings, with strong bootstrap support for major nodes. North American species such as *M. angustifolia*, *M. ioensis*, and *M. coronaria* formed a well-supported monophyletic clade, indicating a shared genetic background. *M. fusca* also formed a distinct group, though some accessions appeared slightly divergent within the clade. *M. florentina* clustered near *M. fusca*, consistent with PCA results. Eurasian wild species, *M. sieversii*, *M. sylvestris*, and *M. orientalis* were grouped closely with *M. domestica*, reflecting their documented contribution to the domesticated apple genome. Several *M. baccata* accessions were interspersed with *M. robusta* and *M. sargentii*, suggesting admixture. The clustering pattern supports clear phylogenetic separation between North American and Eurasian wild species, while also highlighting areas of potential introgression or shared ancestry, particularly among East Asian species and cultivated apples.

**Figure 5:**
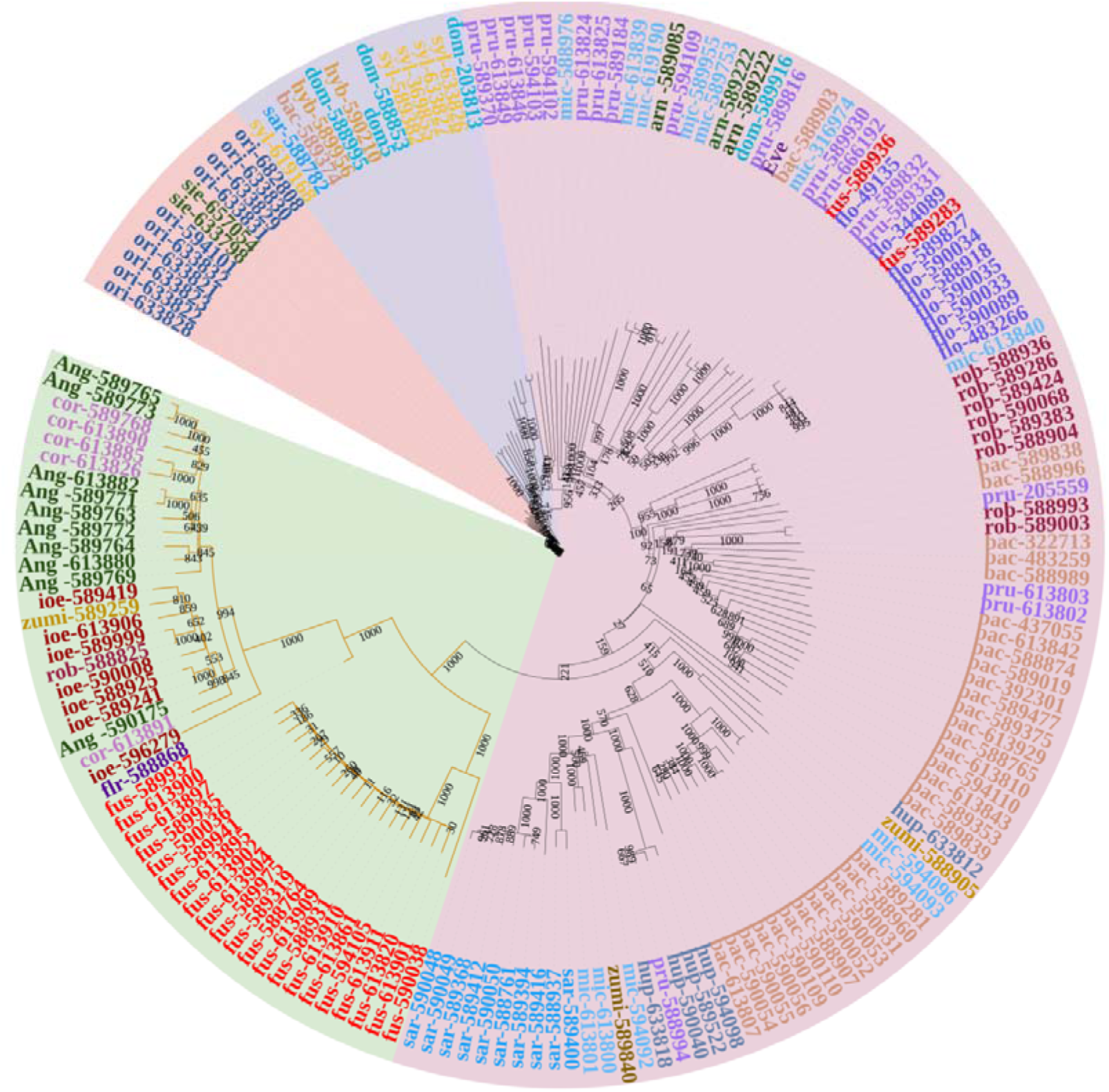
Maximum likelihood phylogenetic tree of diverse *Malus* accessions based on genome-wide SNP data, with accessions color-coded by species: North American species (green), East Asian species (lavender), followed by *Malus sylvestris* from Europe and domesticated *M. domestica* (light blue), and finally *M. sieversii* and *M. orientalis* from Central Asia (peach); bootstrap support values are shown at major nodes. Abbreviations: Ang *= M. angustifolia*, cor = *M. coronaria,* fus *= M. fusca,* ioe *= M. ioensis,* zumi *= M. zumi,* flo *= M. florentina,* bac *= M. baccata,* arn = *M.arnoldiana,* flr *= M. floribunda,* mic *= M. micromalus,* sar *= M. sargentii,* hup *= M. hupehensis,* pru *= M. prunifolia,* rob *= M. robusta,* eve *= M. evereste,* ori *= M. orientalis,* hyb *= M. hyb,* sie *= M. sieversii,* syl *= M. sylvestris,* dom *= M. domestica* and Dom5 =T1190.

#### 2.4 Population structure analysis

Structure analysis, supported by the Evanno ΔK method (Supplementary figure 1), revealed nine distinct populations best represent the ancestry of the *Malus* accessions under study. *M. ioensis, M. coronaria*, and *M. angustifolia* share a common ancestral background, indicating close genetic relatedness among these North American species.

*M. fusca* exhibits a distinct ancestral background, separate from other North American *Malus* species. Two *M. fusca* accessions that did not cluster with the main *fusca* group, did not show the same ancestral background as other *fusca* accessions, instead they, show evidence of admixture with *M. sargentii* and *M. sieverssii*. *M. baccata* represents a separate ancestral population, although many of its accessions exhibit admixture, likely resulting from gene flow with other wild *Malus* species. *M. floribunda* also shows signs of admixture, particularly from other East Asian *Malus* species (Figure 6).

**Figure 6:**
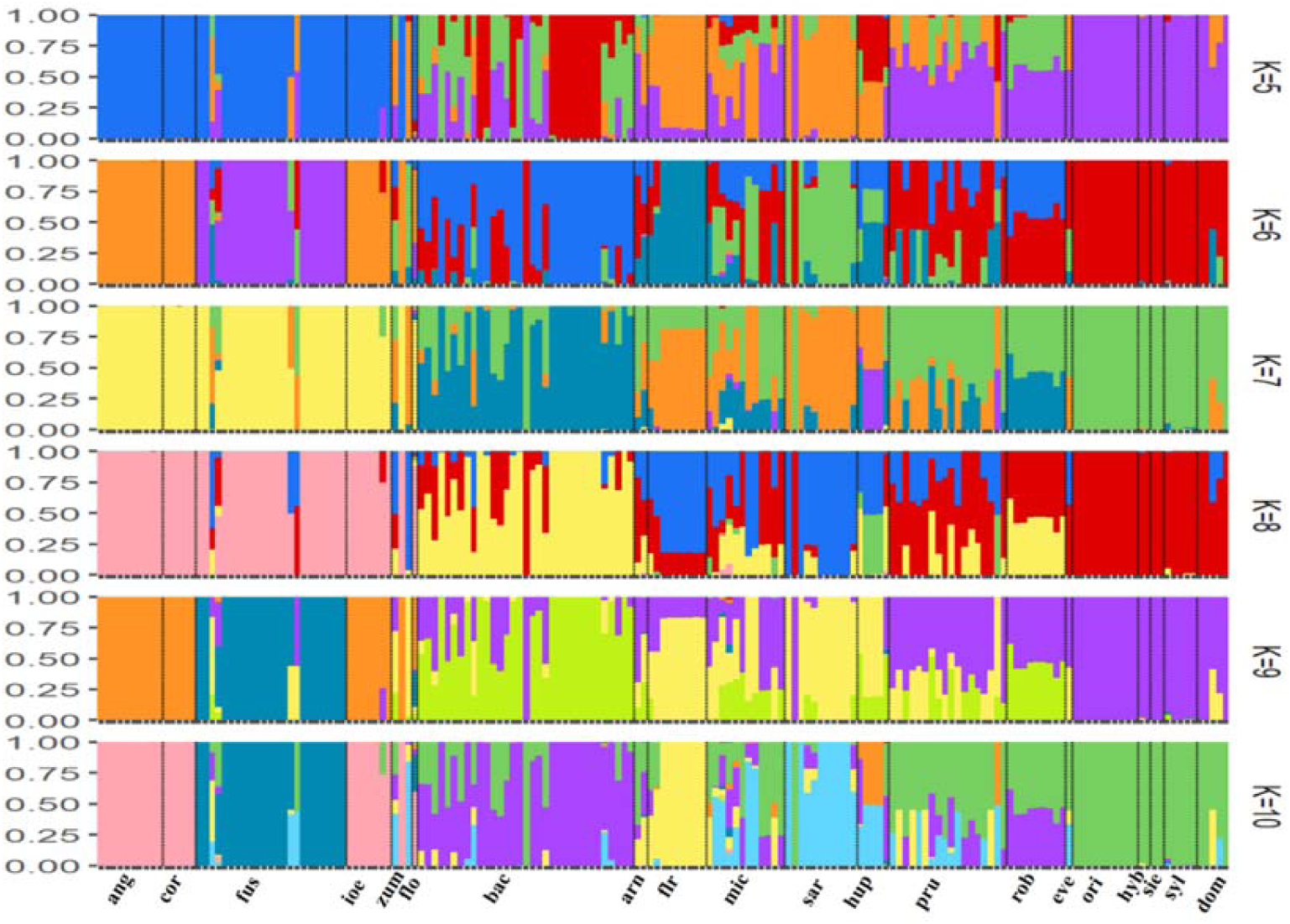
Population structure analysis of *Malus* accessions inferred using STRUCTURE for K = 5 to K = 10. Each vertical bar represents an individual accession, and the colors indicate estimated membership proportions in inferred ancestral populations. Species codes are shown along the x-axis. As K increases, substructure within species becomes more apparent. Abbreviations: ang *= M. angustifolia*, cor = *M. coronaria,* fus *= M. fusca,* ioe *= M. ioensis,* zum *= M. zumi,* flo *= M. florentina,* bac *= M. baccata,* arn = *M.arnoldiana,* flr *= M. floribunda,* mic *= M. micromalus,* sar *= M. sargentii,* hup *= M. hupehensis,* pru *= M. prunifolia,* rob *= M. robusta,* eve *= M. evereste,* ori *= M. orientalis,* hyb *= M. hyb,* sie *= M. sieversii,* syl *= M. sylvestris* and dom *= M. domestica*.

Across the *Malus* diversity population, micro-haplotype analysis revealed substantial variation in the number of haplotypes per locus. Most loci exhibited moderate haplotype diversity, with the number of distinct haplotypes per locus ranging from 1 to over 40. The distribution was right-skewed, with a peak around 9 haplotypes, indicating that the majority of loci (over 300) contained 8–10 haplotypes. A smaller proportion of loci showed high haplotype richness (>20 haplotypes), suggesting regions of elevated allelic diversity or recombination. These patterns reflect a high level of genetic diversity within the panel and support the utility of multi-allelic loci for resolving fine-scale population structure and evolutionary relationships in *Malus* (Figure 7).

**Figure 7:**
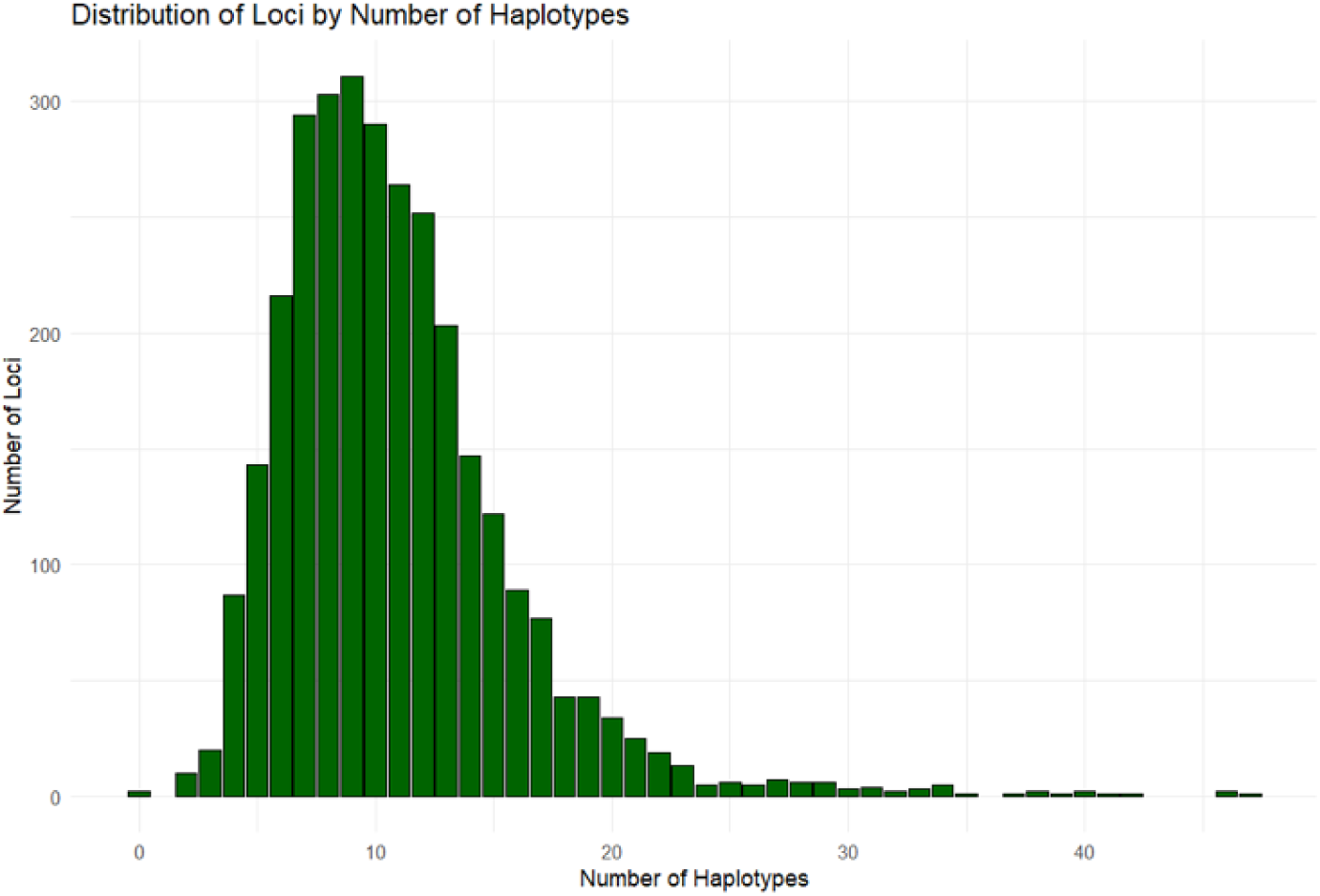
Distribution of loci by number of haplotypes in the *Malus* diversity panel, showing the frequency of loci with varying levels of haplotype diversity across accessions; the x-axis represents the number of distinct haplotypes per locus, while the y-axis indicates the number of loci exhibiting each haplotype count.

#### 2.5 LD decay analysis and GWAS

There is a sharp decline in LD over short distances, with r^2^ values dropping rapidly within the first few kilobases. The half-decay point occurs at approximately 8,000 bp (Figure 8). Genome-wide association studies (GWAS) were conducted to identify SNPs associated with fruit length, weight, and width in an apple diversity panel. All three traits exhibited right-skewed distributions (Supplementary figure 2), with most accessions clustering in the lower value ranges and a smaller number of individuals showing substantially larger fruit dimensions (Supplementary Table 3). The genome- wide association analysis revealed a polygenic basis for fruit size traits, with distinct but partially overlapping loci contributing to variation in fruit length, weight, and width. For fruit length, most accessions measured between 10 and 30 mm, with a few exceeding 60 mm, and significant associations were identified on chromosomes 1, 4, and 9, where several markers surpassed the Bonferroni threshold (–log[J[J(p) = 5), along with moderate associations on chromosomes 5, 6, and 11. The strongest GLM associations for fruit length were observed at *S01_31069925*, *S04_30479303*, and *S09_5414418*, with –log[J[J(p) values of 8.15, 9.58, and 7.67, respectively. In addition to these major loci on chromosomes 1, 4, and 9, several SNPs on chromosomes 5, 6, 11, and 15—such as *S06_31198297*, *S11_41050955*, and *S15_31363354*—showed moderate significance. Fruit weight was skewed toward smaller sizes, with most accessions weighing less than 150 g and relatively few exceeding 200 g. Genome-wide association analysis revealed a highly significant locus on chromosome 4 at *S04_30479303*, with a peak –logCC(p) of 11.86 in the GLM model and 10.48 in the MLM model, indicating a strong and consistent association across models. Additional significant associations were observed at *S09_1227048* (–logCC(p) = 7.13 GLM; 5.86 MLM), *S10_38529651* (–logCC(p) = 6.91 GLM; 6.69 MLM), *S11_41163261* (–logCC(p) = 6.83 GLM; 6.16 MLM), and *S12_31337187* (–logCC(p) = 6.27 GLM; 5.69 MLM). Additional moderate associations included *S05_16541510* (–logCC(p) = 5.88), *S06_31198297* (–logCC(p) = 5.97), and *S01_31810145* (–logCC(p) = 5.58 GLM; 5.55 MLM). The identification of overlapping loci across models supports a polygenic basis for fruit weight, with several loci displaying robust and replicable signals. Among the top associated loci, S01_31810145, S04_30479303, S06_31198297, S09_1227048, S11_41163261, and S12_31337187 were common to both fruit weight and fruit length, indicating shared genetic regulation of these traits. Similarly, fruit width was concentrated between 10 and 30 mm, with some individuals exceeding 75 mm. The strongest association was detected on chromosome 4 at *S04_30479303* (–logCC(p) = 8.97), followed by a moderate association on chromosome 6 at *S06_31198297* (–logCC(p) = 5.95). Both loci were also associated with fruit length and weight, highlighting a shared yet complex genetic architecture underlying fruit size traits in *Malus* (Figure 9) (Supplementary Table 4). Overall, these GWAS results highlight overlapping as well as trait-specific genomic regions associated with apple fruit morphology. The identified loci provide promising targets for further validation and marker-assisted breeding to improve fruit size and shape.

**Figure 8:**
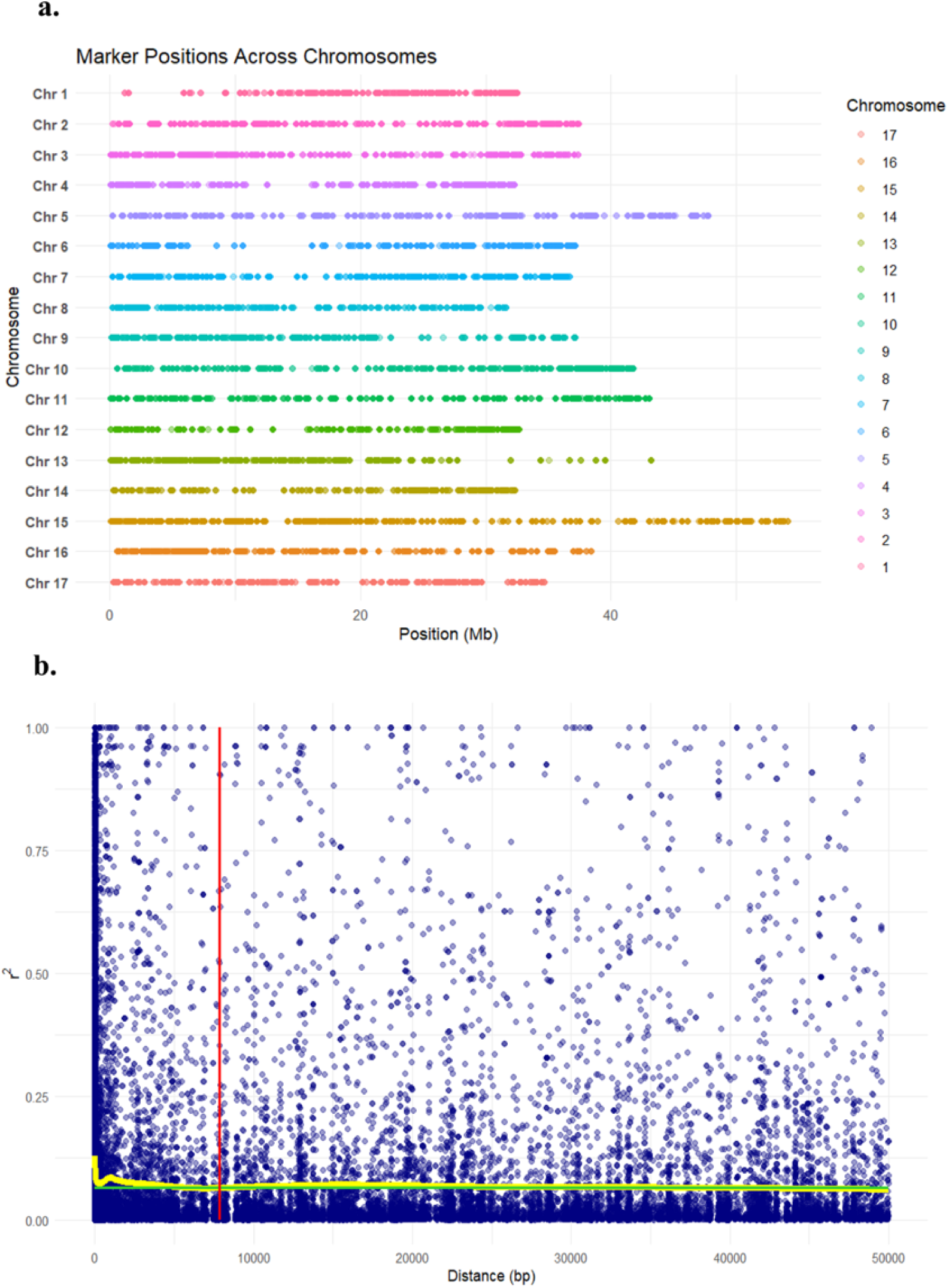
Genomic distribution of markers and linkage disequilibrium decay in *Malus*. **a.** Distribution of genetic markers across the 17 *Malus domestica* chromosomes. Each dot represents the physical position of a marker, color-coded by chromosome. Markers are evenly distributed across the genome, with selection based on gene density to ensure uniform coverage **b.** Linkage disequilibrium (LD) decay plot showing pairwise r² values against physical distance (bp). The yellow LOESS curve represents the average LD decay, with r² declining with increasing inter-marker distance. A red vertical line indicates the approximate LD decay threshold used to inform marker spacing in downstream analyses.

**Figure 9:**
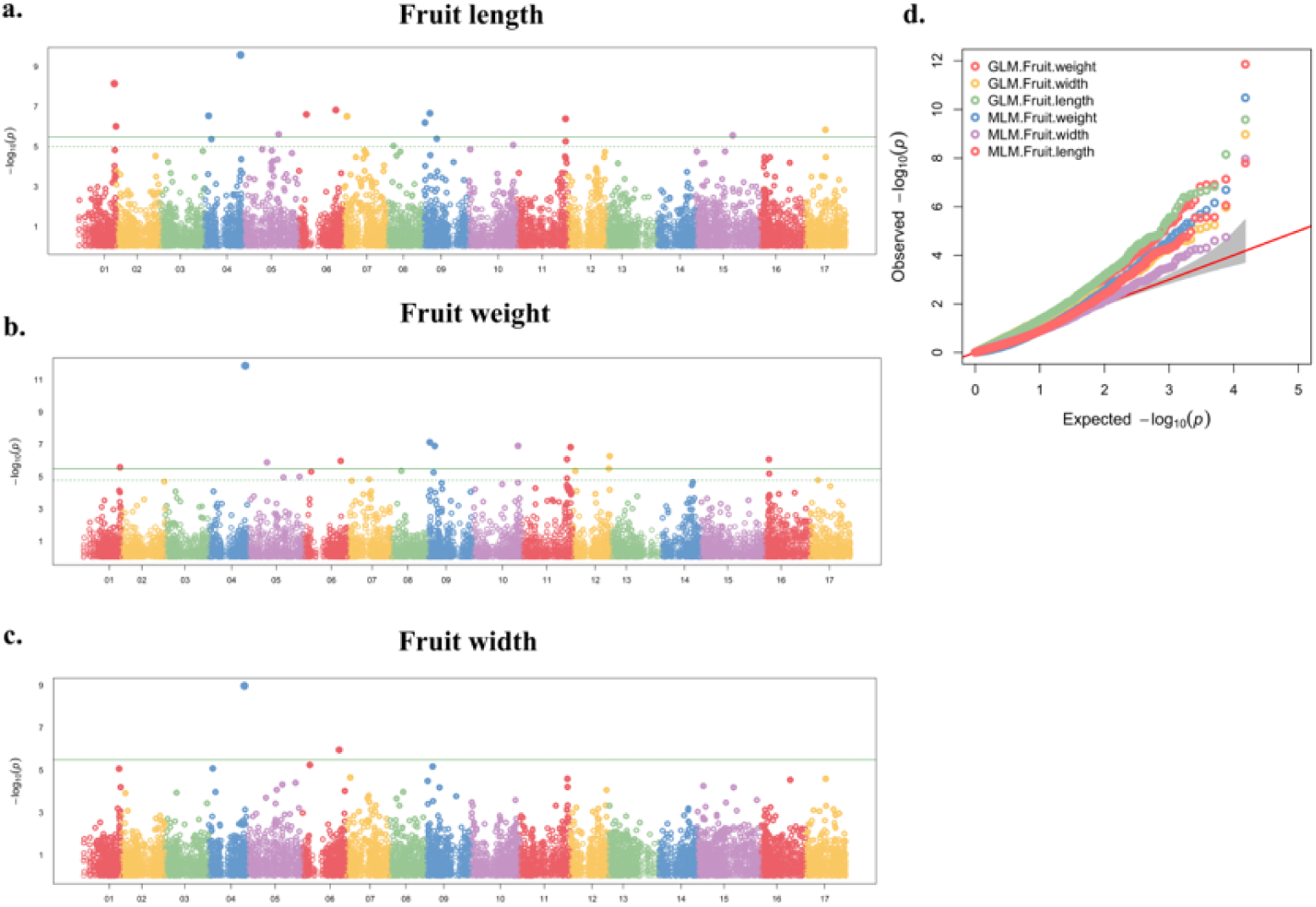
**Genome-wide association analysis for fruit length, weight and width in *Malus.*** Manhattan plots (a–c) display the –logLL(p) values of SNP–trait associations across all 17 chromosomes for: **(a)** fruit length, **(b)** fruit weight, and **(c)** fruit width. The x-axis represents the physical positions of SNPs ordered by chromosome, and the y-axis shows the statistical significance of association (–logLL(p)). Each point represents a single SNP, color-coded by chromosome. The horizontal green dashed line indicates the genome-wide significance threshold (–logLL(p) = 5). **(d)** Quantile– quantile (Q–Q) plot comparing observed vs. expected –logLL(p) values under the null hypothesis for each trait analyzed using both GLM and MLM models. Trait– model combinations are color-coded as indicated in the legend. Deviation from the diagonal red line indicates potential true associations.

## Discussion

Wild relatives of the domesticated apple (*Malus domestica*) represent a valuable reservoir of genetic diversity, particularly for traits related to disease resistance and tolerance to abiotic stresses. This underutilized genepool hold immense potential for accelerating the development of novel cultivars with improved resilience through targeted breeding strategies (Sabety et al. 2024). Here, we present a cost-effective medium density pan-generic SNP and micro-haplotype marker panel optimized to capture genome-wide variation across the *Malus* genus. The marker panel was developed based on genome-wide conserved, syntenic, and collinear genomic blocks identified within the core genome of 13 diverse *Malus* species, thereby enabling cross-species marker transferability across the genus.

*Malus* is a highly heterozygous genus, shaped by extensive gene recombination and natural mutations throughout its evolution. Understanding the genetic architecture underlying valuable traits is essential for advancing fruit tree breeding (Khan et al. 2021). Identifying the molecular mechanisms that drive trait variation is particularly important to meet the increasing demand for high-quality, and sustainably produced pome fruits (Chagne et al. 2012). One of the major challenges in breeding disease resistant apples today is elucidating the complex relationship between phenotypic diversity and underlying genomic variation (Khan et al. 2024; Liu et al. 2022; Švara et al. 2024b). This marker panel was developed and validated to capture genomic variation, including SNPs, INDELs, and micro-haplotypes across the diverse *Malus* genus. While previous studies assessed genetic diversity in wild apple germplasm using a limited number of SSR markers (Zhang et al. 2012; Gao et al. 2015; Wang et al. 2024; Gao et al. 2021), most available SNP arrays were developed using discovery panels composed primarily of *M. domestica* cultivars (Chagné et al. 2012; Bianco et al. 2014, 2016). This domesticated apple–focused nature of the SNP arrays introduces ascertainment bias, resulting in poor representation of rare alleles from wild *Malus* species (Peace et al. 2019; Giebel et al. 2021). Subsequently, the use of the Infinium 20K SNP array for genotyping 178 *Malus sylvestris* accessions revealed that 20% of the markers exhibited more than 10% missing values, 30% had a minor allele frequency (MAF below 5%, and 6% were entirely monomorphic (Buiteveld et al. 2021). In contrast, the DArTag panel we developed has 16,000 universal genetic markers showing less than 0.5% missing data, demonstrating cross-species transferability and applicability across diverse *Malus* accessions.

DArT-based marker panels have demonstrated superior performance over GBS approaches in highly heterozygous and polyploid crops such as alfalfa, strawberry, and sweetpotato, offering increased genome coverage, reduced missing data, improved genotyping accuracy, and enhanced cross-species transferability (Zhao et al. 2023; Zhao et al. 2024; Jose et al. 2015). DArTag technology addresses this need by designing oligonucleotides that target known SNPs and small INDELs (<50 bp), along with their flanking regions. The resulting amplicons, 54 bp or 81 bp are sequenced using next-generation sequencing (NGS), allowing for the detection of multiple alleles per locus (Zhao et al. 2023 & 2024). Unlike traditional SNP arrays or KASP assays, which typically detect only two alleles per site, DArT enables identification of short multiallelic haploblocks (Zou et al. 2020; Jose et al. 2015). In this study, the DArT markers exhibited an average of 9-10 alleles per locus across the *Malus* diversity panel, highlighting their high resolution and informativeness. Similar results have been observed in sweetpotato, where 183 marker loci revealed more than 10 distinct micro-haplotypes, reflecting extensive genetic diversity. Compared to biallelic SNPs, these multiallelic haplotype markers provide greater information content, simplify haplotype phasing, and improve inference of ancestry. The even genomic coverage and reduced ascertainment bias associated with DArTag marker panel makes it particularly valuable for genetic studies in highly heterozygous systems, as well as for applications in population genetics and ecological genomics (Zhao et al. 2023; Zhao et al. 2024).

This pan-generic marker panel offers high-resolution, gene-informed marker distribution, suitable for large-scale genotyping, QTL validation, and genome-wide association studies across diverse *Malus* populations. To ensure uniform genome- wide coverage, marker selection was refined based on gene density resulting in the retention of 3,100 gene-informed genomic blocks including 104 SNPs linked to previously identified important QTLs. This design strategy provides balanced representation of both coding and regulatory regions, enhancing the utility of the panel for functional and evolutionary genomics. Notably, markers exhibiting high allele frequency variance between wild and domesticated apples were prioritized, making them particularly informative for studying hybrid populations and introgression patterns. Additionally, markers with low F_st_ but high MAF between wild and cultivated groups offer valuable resolution for capturing shared genetic diversity across *Malus*, increasing their applicability in broad population genomic analyses. The panel enables fine-scale marker–trait association discovery, dissection of the genetic architecture of complex traits, and the study of both neutral and adaptive genetic variation. To validate the performance of the panel, it was applied to three bi- parental mapping populations and a diverse *Malus* germplasm panel. Across these datasets, more than 1,500 SNP and haplotype markers were identified as informative, underscoring the panel’s robustness and versatility. PCA further demonstrated its discriminatory power, with FC individuals clearly segregating between both parents. This pattern confirms the panel’s high resolution and reliability in capturing inherited genetic variation and population structure. Similar results have been found in DArT panels developed for potato (*Solanum tuberosum L.*) (Endelman et al. 2024), strawberry (*F. × ananassa*) (Hardigan et al. 2023), and alfalfa (Zhao et al. 2023). The 3K DArTag panel for apple offers substantial practical benefits, particularly in terms of scalability and cost-efficiency. With a per-sample cost of less than $15, it provides an affordable solution for high-throughput genotyping in large-scale studies. This makes it especially suitable for bi-parental mapping populations, where it enables the construction of high-resolution linkage maps while simultaneously allowing the detection of additional loci associated with important agronomic traits. Its high informativeness, genome-wide coverage, and low ascertainment bias make it very attractive for genetic studies in complex, heterozygous systems such as *Malus*.

The pan-generic marker panel was validated on a broad *Malus* diversity panel, revealing clear phylogenetic structure and supporting the panel’s resolution in detecting both broad-scale divergence and fine-scale variation within and between species. Phylogenetic clustering identified three major clades. Clade I comprised of North American *Malus* species, including *M. ioensis*, *M. coronaria*, *M. angustifolia*, and *M. fusca*. Within this clade, *M. angustifolia* and *M. coronaria* formed a close sister group, while *M. ioensis* clustered as a more distinct subgroup. Interestingly, one accession of *M. coronaria* (PI 613891) collected from Ontario and a *M. angustifolia* accession PI 590175, collected from New York clustered with *M. ioensis*, whereas the remaining *M. coronaria* accessions grouped with *M. angustifolia*. These clustering patterns may reflect admixture zones, historical gene flow within North American *Malus*, or possible misclassification. *Malus fusca*, native to the Pacific Northwest and commonly referred to as the Oregon or Pacific crabapple, formed a genetically distinct group, in agreement with recent pan-genome studies that place *M. fusca* as divergent from other North American species (Svara, 2024a; Li et al. 2024). Notably, two accessions of *M. fusca* (PI 589941 and PI 590038) from California and one from an unspecified U.S. location clustered apart from the main *M. fusca* group, suggesting mis-labelling or potential admixture.

A second major cluster consisted of East Asian *Malus* species, including *M. arnoldiana*, *M. baccata*, *M.* ‘Evereste’, *M. floribunda*, *M. hupehensis*, *M. micromalus*, *M. prunifolia*, *M. × robusta*, *M. sargentii*, and *M. zumi*. A third clade comprised domesticated apples (*M. domestica*) and its progenitor species including *M. sieversii*, *M. sylvestris*, and *M. orientalis*. This grouping reflects known evolutionary histories, where *M. domestica* originated in Central Asia from *M. sieversii*, with introgression from *M. sylvestris* and *M. orientalis* during the process of domestication and westward dispersal (Duan et al. 2017).

These findings underscore the capacity of this pan-generic marker panel to detect genetic relationships across the *Malus* genus. The clear separation of major geographic groups, detection of intra-specific variation, and identification of potential admixture or mislabeling events highlight its utility for taxonomy, germplasm curation, evolutionary biology, and pre-breeding applications. Moreover, the panel’s effectiveness in resolving complex polyploid and hybrid structures provides a strong foundation for its use in trait mapping, genomic selection, and conservation strategies across wild and cultivated *Malus* populations.

We identified multiple loci significantly (–logCC(p) ≥ 5) associated with fruit length, weight, and width, highlighting the polygenic and partially overlapping genetic architecture of fruit size traits. The most robust associations were observed on chromosomes 1, 4, 5, 6, 9, 11, 12, and 15, with *S04_30479303* and *S06_31198297* consistently detected across all traits. The detection of major loci on chromosomes 4, 5, 6, 9, 11, and 15 is broadly consistent with the results reported by Devoghalaere et al. (2012), who identified six QTLs for fruit weight on LGs 5, 8, 11, 15, 16, and 17 using bi-parental populations derived from ‘Royal Gala’ × ‘Braeburn’ and ‘Starkrimson’ × ‘Granny Smith’. Their study reported conserved QTLs on LGs 8 and 15, along with epistatic interactions among loci on LGs 8, 15, and 17. In line with these findings, significant associations in the current study were also observed on chromosomes 5, 11, and 16, with *S05_16541510* and *S16_3804813* influencing fruit weight, and *S11_41050955* and *S11_41163261* associated with both fruit weight and length. The repeated detection of signals on chromosome 15 across independent studies highlights its potential role as a key genomic region involved in the regulation of fruit size in apple. Moreover, our findings are in strong agreement with the genome-wide QTL analysis conducted by Chang et al. (2014), who used a ‘Jonathan’ × ‘Golden Delicious’ population to identify multiple QTLs for fruit size, shape, length, and diameter across at least 11 linkage groups. Their results highlighted LGs 4, 5, 8, 11, 12, 15, and 17 as recurrent hotspots for fruit-related traits. Notably, QTLs for both fruit length and diameter were mapped to LGs 4 and 11. Our study similarly detected significant (–logCC(p) ≥ 5) associations for fruit size traits on chromosomes 4, 5, and 11, and we additionally identified *S04_30479303* on chromosome 4 as a major shared locus across all three traits, supporting the hypothesis that this region harbors genes of broad regulatory influence.

Overall, our study confirms the highly polygenic and partially overlapping genetic architecture underlying fruit morphology in apple. Importantly, the ability of the DArTag panel to detect both major and minor associations across a diverse panel confirms its effectiveness not only for diversity analysis but also for high-resolution GWAS applications in apple.

## Conclusion

We report the successful development of a pan-generic 3K DArTag marker panel tailored for comprehensive genomic applications in *Malus* species. This cost-effective, high-throughput, transferable marker system enables robust QTL mapping, GWAS, linkage map construction, and population genomics analyses across diverse genetic backgrounds. The panel’s broad cross-species utility and genome-wide coverage provide a valuable resource for dissecting trait architecture, facilitating marker- assisted selection, and accelerating breeding efforts in apples and its wild relatives.

## Declarations Availability of data

All data used in the manuscript is cited or included in the manuscript.

## Ethics approval and consent to participate

Not applicable

## Consent for publication

Not applicable

## Acknowledgments

We acknowledge Della Cobb-Smith for her help to collect leaf samples for DNA extraction and genotyping.

## Author Contributions

A.Kh conceived the study, acquired the funding, and supervised the research. R.T, H.F, A.Š, and P.C, collected the leaf samples for DNA extraction. A.S. performed bioinformatic analysis, conducted genetic analysis, and wrote the original draft of the manuscript. J.G and Q.S, performed bioinformatic analysis for the oligo design. A.S, A.Ki and A.Kh, revised the manuscript. All authors have read and approved the manuscript.

---

**Supplementary Figure 1:**
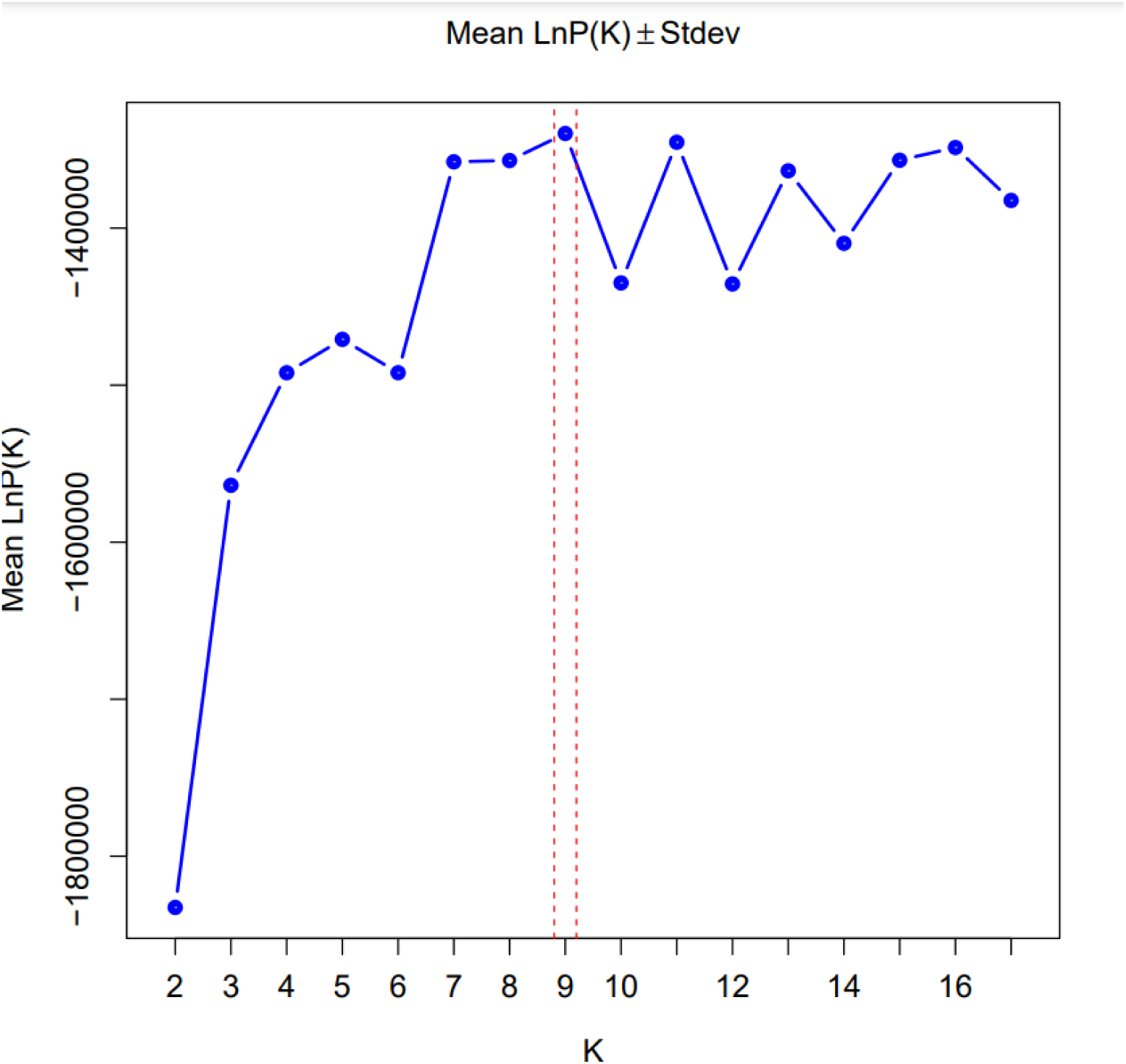
Evanno plot showing ΔK values for determining the optimal number of genetic clusters.Mean log probability of the data [LnP(K)] ± standard deviation is plotted against the number of clusters (K). Vertical red lines indicate the optimal K as determined by the Evanno ΔK method, supporting the presence of nine genetically distinct populations in the dataset.

**Supplementary Figure 2:**
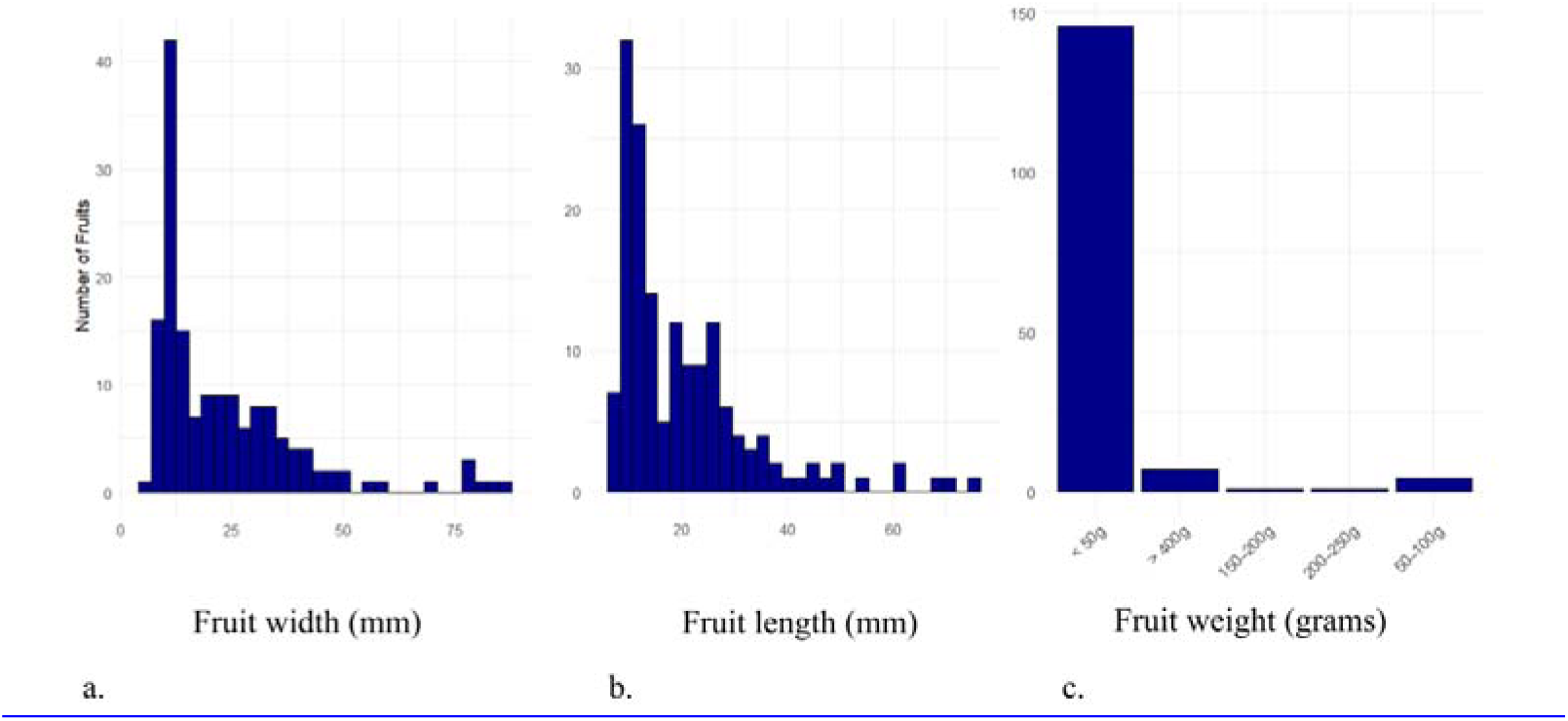
Distribution of fruit morphological traits across *Malus* **accessions.** Histograms show the frequency of fruits measured for a. fruit width, b. fruit length, and c. fruit weight. Width and length are shown in centimeters, with most fruits clustering below 40 cm in both dimensions. Fruit weight is categorized into discrete bins:L<50Lg, 50–100Lg, 150–200Lg, 200–250Lg, and >400Lg, with the majority of fruits weighing less than 50Lg. The y-axis represents the number of fruits observed within each bin.

